# Berberine, the major bioactive compound from *Berberis aristata,* attenuates virulence of multidrug resistant *Chromobacterium violaceum* at non-lethal concentrations by targeting bacterial efflux and denitrification machinery

**DOI:** 10.1101/2025.10.17.683191

**Authors:** Nidhi Thakkar, Gemini Gajera, Vijay Kothari

**Affiliations:** Institute of Science, Nirma University, Ahmedabad, India

**Keywords:** Antimicrobial Resistance (AMR), *Caenorhabditis elegans*, Denitrification, Emerging Pathogen, Transcriptome, Virulence

## Abstract

*Berberis aristata* root extract, and berberine were assessed for their anti-pathogenic activity against multidrug resistant *Chromobacterium violaceum*. Berberine was found to be more potent than the parent extract with respect to attenuation of the pathogen’s virulence against the model host *Caenorhabditis elegans.* It also performed better than six different antibiotics, when compared at same concentration. Berberine was able to modulate multiple traits of the target pathogen, such as, biofilm formation, haemolytic activity, efflux/transport activity, and exopolysaccharide production. Repeated exposure of *C. violaceum* to berberine did not induce resistance. Whole transcriptome analysis of the berberine-treated *C. violaceum* revealed differential expression of genes associated with stress response, efflux, denitrification, and metalloproteases. Downregulation of two genes of the denitrification pathway, *nirK* and *norB*, was confirmed through RT-PCR too.

## 1. Introduction

Bacterial pathogens have remained among the biggest challenges to human health and welfare since antiquity. New pathogens, and new antibiotic-resistance patterns keep emerging every few decades. While development of novel antibacterial solutions is urgently required against Priority Pathogens listed by the World Health Organization (WHO; (https://www.who.int/publications/i/item/9789240093461) and the Centers for Disease Control and Prevention (CDC; https://www.cdc.gov/antimicrobial-resistance/data-research/threats/index.html), keeping an eye on emerging pathogens is also important in order to be better prepared for possible future outbreaks. This study focuses on the bacterium *Chromobacterium violaceum*, which is being viewed as a pathogen of emerging importance (Batista and da Silva Neto, 2017; Kothari et al., 2017). While multidrug resistant (MDR) strains of *C. violaceum* are recognized as serious threat (Pei et al., 2024), this bacterium is also considered a good model to study quorum sensing (QS) in bacteria. It is noteworthy that QS, a mechanism of intercellular communication among bacteria, is also viewed as a plausible target in pathogenic bacteria (Revanasiddappa et al., 2024). Owing to the production of the violet bioactive pigment, violacein, in *C. violaceum* being regulated by QS, and inherent resistance to multiple antibiotics, it has become an important model organism for researchers working in the area of antimicrobial development.

*C. violaceum* is a gram-negative, facultative anaerobic, non-sporulating, motile, beta proteobacterium, which though does not infect large number of people, mortality rate among patients infected by this bacterium is high (20 to 60%). Virulence factors of this pathogen include violacein pigment, hydrogen cyanide (HCN), biofilm formation, Type III Secretion System (T3SS) effector proteins, etc. (Venkatramanan and Nalini, 2024). This pathogen can cause severe infections such as skin lesions, sepsis, and liver abscesses, particularly in immunocompromised individuals. Multiple cases of *C. violaceum* infection have been reported from different countries e.g. United States, Cuba, Australia, Japan, Malaysia, Brazil, Vietnam, Sri Lanka, Taiwan, Singapore, Argentina, Nigeria, Canada, and India. This organism is part of the natural microbiota of water and soil. The most prominent way for it to enter the bloodstream and cause systemic infections is through wounds or cuts in the skin, where the bacterium enters from a polluted surface or water (Alisjahbana et al., 2021).

Novel anti-pathogenic approaches, not necessarily antibiotic-based, are urgently required against different multidrug resistant pathogenic bacteria including *C. violaceum*. The ‘anti-virulence’ strategy is one such non-antibiotic approach, wherein the ‘pathoblockers’ are expected to attenuate bacterial virulence with no or little effect on its growth. Many plants, polyherbal formulations from traditional medicine, and certain phytocompounds have been reported for their anti-virulence mode of action against pathogenic bacteria (Patel et al., 2020a). Such virulence-targeting formulations are expected to exert lesser selection pressure on susceptible pathogens to develop resistance. *Berberis aristata* is one of the widely prescribed plant in the Indian system of traditional medicine, the *Ayurved*. The plant is used to treat diabetes, allergies, conjunctival inflammation and infection, ulcers, and as laxative, tonic, and blood-purifying agent. Among other indications for the use of *B. aristata*, frequently mentioned, are liver/gallbladder/pancreatic disorders, difficult-to-heal wounds and skin inflammation or infection (ulcers, abrasions), hemorrhoids, malaria, leprosy, dysentery, urinary tract ailments, plague infection and menopause (Marchelak et al., 2025). One of the major phytocompounds from this plant is berberine, which is an isoquinoline alkaloid. Though both *B. aristata* and berberine have been reported to possess antimicrobial activity, many such reports have limited themselves to qualitative disc diffusion assays (Saxena et al., 2014), or have shown the activity at ‘mg’ levels (Thakur et al., 2020). Detailed investigation, which are more quantitative and offer mechanistic details associated with the antibacterial activity of this plant or berberine are warranted. In this context, the present study investigated anti-pathogenic potential of *B. aristata* and berberine against *C. violaceum* through *in vitro*, *in vivo*, and transcriptome analysis.

## 2. Methods

### 2.1. Test plant and compound

Powder of the *Berberis aristata* DC. (Family Berberidaceae) roots was procured from Yucca Enterprises, Mumbai. This plant sample was divided into two parts. One part was dissolved in dimethylsulfoxide (DMSO; Merck), and the other one was subjected to hydroalcoholic extraction. Three grams of the plant powder was suspended into 8 mL of DMSO, and subjected to shaking at 100 rpm for 30 minutes at room temperature. This was followed by centrifugation at 8000 rpm for 20 minutes. The supernatant obtained was used as DMSO-soluble fraction of *B. aristata* roots. Solubility of the plant power in DMSO was calculated to be 9.87%.

For preparing hydroalcoholic extract of the *B. aristata* roots, 5 g of the root powder was suspended into 50 mL of the solvent (water: ethanol in 1:1 ratio), and subjected to shaking at 100 rpm for 2 h at ambient temperature. The resulting liquid was centrifuged at 8000 rpm for 20 min. Insoluble fraction obtained in the pellet form was discarded. The supernatant was dried, followed by reconstitution in DMSO. Extraction efficiency, calculated as percentage weight of the starting dried plant material, was found to be 3.08%. Reconstitution efficiency (i.e. solubility of the dried hydroalcoholic extract in DMSO) was 89.75%.

Both, DMSO-soluble fraction of *B. aristata* roots (DMSO-BAR), and hydroalcoholic extract of the roots (HA-BAR) reconstituted in DMSO were stored in sterile glass vials (15 ml, Merck) under refrigeration (4-8°C). Inner surface of caps of these vials was covered with aluminium foil to prevent possible leaching of the cap material into the solvent.

For preparing berberine for assay purpose, 20 mg of berberine chloride (Sigma-Aldrich) was dissolved in 1 mL DMSO.

### 2.2. Test organisms

#### Pathogen

*C. violaceum* strain (MTCC 2656) was procured from Microbial Type Culture Collection, Chandigarh. Its antibiogram (Table S1) generated through disc diffusion assay revealed it to be resistant to three different classes of antibiotics, that is, Polymyxin (colistin), Lincosamide (clindamycin), and β-Lactam (cefixime, cefpodoxime, and augmentin). Nutrient broth or agar (HiMedia) was used to cultivate this bacterium. Inoculum density of this bacterium to be used in all experiments was adjusted at OD_625_=0.08-0.10 to achieve equivalence to McFarland turbidity standard 0.5.

#### Model host

The nematode worm *Caenorhabditis elegans* (N2 Bristol), procured from the Caenorhabditis Genetics Center (USA) was used as a model host for *C. violaceum*. Worm synchronization was done as described in literature (Corsi et al., 2015), and in our previous studies (Joshi et al., 2019; Patel et al., 2019a) too. Lyophilized *Escherichia coli* OP50 procured from Biovirid (The Netherlands) was used as food for *C. elegans*, while maintaining the worm on NGM (nematode growth medium) agar. Prior to all *in vivo* assays, worms were kept in M9 buffer without food for 2 days to make them gnotobiotic.

### 2.3. *In vivo* assays

Four different types of *in vivo* assays (Parmar et al., 2024) performed are described in the following text, wherein the nematode *C. elegans* was used as a model host for the bacterial pathogen *C. violaceum*. All these assays involved live-dead counting of worm population over a five-day period, under a light microscope (4X) equipped with halogen light source. On final day of the experiment, when plates could be opened, death of worms was confirmed by touching them with a straight wire, wherein absence of movement was considered as confirmation of death. These assays were performed in 24-well plates (surface non-treated; HiMedia). Each well contained ten gnotobiotic worms (L3-L4) in M9 buffer, who were challenged with *C. violaceum* by adding 100 µL (OD_764_=1.50 ± 0.05) of bacterial culture grown in nutrient broth for 22 ± 1 h at 35 ± 0.5°C. Total assay volume in each well was 1 mL.

#### 2.3.1. Anti-pathogenic assay

Ten worms (L3-L4 stage) contained in 900 µL of M9 buffer were challenged with *C. violaceum* (100 µL of the culture broth) in absence or presence of the test extract/berberine, wherein neither the bacterium nor the worms were pre-exposed to the plant product. Incubation was done at 22°C for 5 days with live-dead microscopic count once everyday.

#### 2.3.2. Anti-infective assay

*C. violaceum* was grown at 35°C for 22-24 hours in nutrient broth with or without the plant product. Post-incubation, 100 µL of the culture broth was mixed with 900 µL of M9 buffer containing ten worms (L3-L4) in a 24-well plate. This plate was incubated at 22°C for 5 days, and worm survival was quantified on daily basis.

#### 2.3.3. Post-infection assay

Ten worms contained in M9 buffer were first challenged with pathogen, and after allowing *C. violaceum* (100 µL of the culture broth) for 24 or 48 hours to establish infection, berberine (10 µg/mL) was added into the well as a possible post-infection therapy. Survival of worms was observed through a live-dead microscopic count over 5 days (Patel et al., 2019b).

#### 2.3.4. Repeated exposure

To test whether repeated exposure of the test pathogen to berberine can induce any resistance in it, we subcultured *C. violaceum* in berberine (10 or 20 µg/mL)-containing broth for ten times, and then the ‘berberine-habituated’ culture thus obtained was tested for its virulence towards the nematode host, in absence or presence of berberine.

Appropriate controls were included in all the above experiments as relevant:

Sterility Control: Sterile M9 buffer containing neither worms nor bacteria

Survival Control: M9 buffer containing ten worms (no bacteria added)

Toxicity Control: Ten worms in M9 buffer supplemented with plant product

Infection Control: Ten worms in M9 buffer + 100 µL of the *C. violaceum* culture broth (OD_764_ = 1.50 ± 0.05). No plant extract was added in these wells.

Vehicle Control: Ten worms in M9 buffer + 100 µL of the *C. violaceum* culture broth (OD_764_ = 1.50 ± 0.05) + DMSO (0.5% v/v) in place of plant product.

Positive Control: Standard antibiotics employed as positive controls are mentioned in the figure legends.

### 2.4. *In vitro* assays

#### 2.4.1. Growth and pigment quantification

The broth dilution assay was used to evaluate *C. violaceum*’s growth and quorum sensing (QS)-regulated pigment synthesis in the presence or absence of the test extract/compound. Different concentrations (5-100 μg/mL) of *Berberis aristata* root extract or berberine were used to challenge the organism. The growth media employed was nutrient broth, into which bacterial inoculum set to 0.5 McFarland turbidity standard was added at 10% v/v, followed by incubation for 22-24 hours at 35° C, with intermittent shaking. The assay system also included an appropriate vehicle control with DMSO (0.5% v/v) and an abiotic control with extract/berberine and growth medium but no bacterial inoculum.

Bacterial cell density was quantified photometrically at the end of the incubation at 764 nm (Agilent Cary 60 UV-visible spectrophotometer) (Joshi et al., 2016). Following this, for extraction and quantification of the violet pigment violacein, one mL of culture broth was centrifuged (13,600 g) for 10 minutes. Cell pellet was dissolved in 1 mL of DMSO (Merck). This allowed the pigment to be extracted in the DMSO. This was followed by centrifugation (13,600 g) for 10 minutes, and OD_585_ of the supernatant was quantified. Violacein Unit was calculated as OD_585_/OD_764_.

#### 2.4.2. Biofilm assay

Biofilm formation is an important virulence trait, and hence the effect of berberine (10-100 µg/mL) on biofilm forming ability of *C. violaceum* was investigated. In this assay, control and experimental, both groups contained three test tubes. Total assay volume was 2 mL. All these nutrient broth (with or without berberine) tubes were inoculated with *C. violaceum* inoculum (10%v/v), followed by incubation at 35°C for 24 h under static condition, which resulted in formation of biofilm as a ring on walls of the glass tubes. Biofilm quantification was achieved through crystal violet assay (Hirshfield et al., 2008), wherein the biofilm-containing tubes (after discarding the inside liquid) were washed with phosphate-buffer saline (PBS) in order to remove all non-adherent (planktonic) bacteria, and air-dried for 15 minutes. Each of the washed tubes was then stained with 1.5 mL of 0.4% aqueous crystal violet (Central Drug House, Delhi) solution for 30 min. Each tube was then washed twice with 2 mL of sterile distilled water and immediately de-stained with 1.5 mL of ethyl alcohol (95%). After 45 min of de-staining, one mL of de-staining solution was transferred into separate tubes, and read at 580 nm (Agilent Cary 60 UV-Vis).

#### 2.4.3. Exopolysaccharide (EPS) Quantification

*C. violaceum* was grown in 20 mL of nutrient broth contained in 100 mL flasks (containing or not containing berberine (10-100 µg/mL). Incubation was done at 35 ^◦^C for 22 ± 1 h under shaking conditions (120 rpm). Post-incubation, cell density was measured as OD_764_. For EPS quantification (Andhare et al., 2017), the culture broth was centrifuged (13,600 g; 10 min), and to the resulting supernatant, chilled acetone (Merck) was added in a 1:2 ratio, and then allowed to rest for 30 min for the EPS to precipitate. The precipitated EPS was then separated by filtration through a pre-weighed Whatman #1 filter paper (Axiva). Filter paper containing EPS on it was then subjected to drying at 55 ^◦^C for 24 h, and the weight of the EPS on the paper was calculated.

#### 2.4.4. Efflux Assay

An ethidium bromide (EtBr) efflux assay with *C. violaceum* was performed using the method described in Mullin et al. (2004). *C. violaceum* cells, grown overnight in nutrient broth were loaded with 10 µg/mL of EtBr (HiMedia) in the presence of 100 µg/mL of reserpine (HiMedia; positive control) or berberine (10, 50 or 100 µg/mL). An effective efflux inhibitor such as reserpine is expected to inhibit efflux during this loading step. Cells were incubated at 35 ^◦^C for 20 min, followed by pelleting through centrifugation (13,600 g; 10 min). The medium was then decanted, and the cell pellet was resuspended in fresh nutrient broth, either with or without berberine, to an optical density of OD_764_ = 0.20. The EtBr efflux was then determined by measuring fluorescence at excitation and emission wavelengths of 530 and 600 nm respectively in a spectrofluorometer (JASCO FP-6500; Jasco International Co. Ltd., Tokyo).

#### 2.4.5. Hemolysis assay

OD_764_ of C. violaceum culture grown overnight in nutrient broth in presence or absence of berberine (10, 50 or 100 µg/mL) was set to 1.00. Cell free supernatant was then prepared by centrifugation at 13,600 g for 10 min. Ten μL of human blood (sourced from author’s own body in healthy condition under no antibiotic treatment; collected in heparinized vial) was incubated with this cell free supernatant for 2 h at 37 °C, followed by centrifugation at 800 g for 15 min. Supernatant was then read at 540 nm, to quantify the amount of haemoglobin released. 1% Triton X-100 (CDH, New Delhi) was employed as the positive control. Phosphate buffer saline (PBS; pH 6.8) was employed as the negative control.

### 2.5. Whole Transcriptome Analysis

To gain insight into the molecular mechanisms by which berberine attenuates bacterial virulence and modulates various traits like efflux, biofilm, haemolysis, etc., we compared the gene expression profile of berberine pre-treated *C. violaceum* with that of control culture at the whole transcriptome level. The overall workflow of this whole transcriptome analysis (WTA) is given in Figure-S1.

#### 2.5.1. RNA extraction

Trizol (Invitrogen Bioservices; 343909) was used to extract RNA from bacterial cells (Jahn et al., 2008). RNA was dissolved in nuclease-free water after precipitation with isopropanol and washing with 75% ethyl alcohol. Using the RNA HS assay kit (Thermo Fisher; Q32851), the extracted RNA was quantified using a Qubit 4.0 fluorometer (Thermo Fisher; Q33238). RNA concentration and purity were evaluated using Nanodrop 1000. Finally, RNA was checked on the TapeStation using HS RNA ScreenTape (Agilent) to yield RIN (RNA Integrity Number) score (Table S2).

Library preparation: Final libraries were measured using a Qubit 4.0 fluorometer (Thermo Fisher; Q33238), a DNA HS assay kit (Thermo Fisher; Q322851), and a TapeStation 4150 (Agilent) using high-sensitivity D1000 ScreenTapes (Agilent; 5067-5582). The acquired sizes of all libraries are reported in Table S2.

#### 2.5.2. Genome annotation and functional analysis

FastQC v.0.11.9 (default parameters) was used to undertake a quality assessment of the sample’s raw fastq readings (Andrews, 2018). The read’s quality was then reevaluated using Fastq v.0.20.1 (Chen et al., 2018) after pre-processing the raw fastq reads with Fastq v.0.20.1.

The *C. violaceum* (ATCC 12472) genome (GCF_000007705.1) was indexed using bowtie2-build v2.4.5 (default parameters). The processed reads were mapped to the *C. violaceum* genome using bowtie2 v2.4.5. Gene counts were determined using feature count v.2.0.1 to quantify the aligned reads from the individual samples. Differential expression was estimated using the exact test (parameters: dispersion 0.1) with these gene counts as inputs in edgeR (Robinson et al., 2010). The up- and down-regulated sequences were extracted from the *C. violaceum* coding file and annotated using Blast2GO (Conesa and Götz, 2008) to obtain the Gene Ontology (GO) keywords. All the raw sequence data has been submitted to the Sequence Read Archive. The relevant accession number of *C. violaceum* and berberine-exposed *C. violaceum* are SRX21456278 and SRX21456279 respectively.

### 2.6. Reverse Transcription-Polymerase Chain Reaction (RT-PCR)

Polymerase chain reaction (PCR) was used to confirm the differential expression of selected DEG revealed from whole transcriptome analysis (WTA). Primer3Plus (Untergasser et al., 2012) was used to design primers for the target genes (Table 1). These primer sequences were verified for their specific and exclusive binding to the target gene sequence throughout the whole genome file of *C. violaceum*. RNA extraction and purity check was executed as described in the previous section. The SuperScript™ VILO™ cDNA Synthesis Kit (Invitrogen Biosciences) was used to generate complementary DNA. Using gene-specific primers purchased from Sigma-Aldrich, the PCR experiment was carried out employing the time-temperature profile shown in Table S3. The gene CV_RS01880 (Q7P125) was kept as an endogenous control. The reaction mix used was FastStart Essential DNA Green Master mix (Roche; 06402712001). Real-time PCR (RT-PCR) assay was performed on QuantStudio 5 RT-PCR machine (Thermo Fisher Scientific, USA). RT-PCR was done from the same genetic material which was used for WTA, as well as from a sample generated independent of that for transcriptome assay.

**Table 1.**
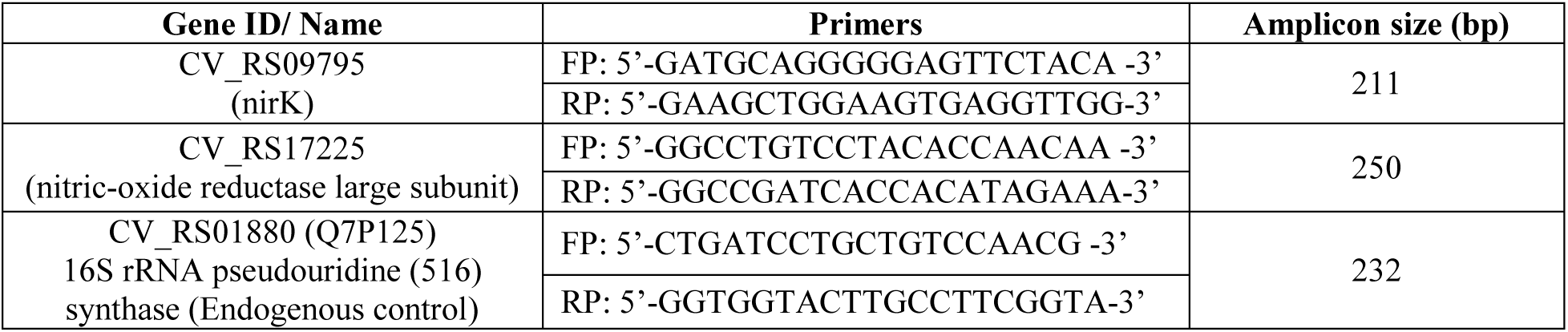
Primer sequences for the target genes.

### 2.7. Statistical analysis

All results reported are means of three or more independent experiments, each performed in triplicate. Statistical significance was assessed through *t*-test performed using Microsoft Excel^®^ (2016), and data with p ≤ 0.05 was considered to be statistically significant.

## 3. Results and Discussion

### 3.1. Neither *B. aristata* nor berberine affected *C. violaceum* growth and Quorum Sensing (QS)-regulated pigment production

To decide appropriate concentration of the plant extract and berberine to be used in further assays, we challenged *C. violaceum* with different concentrations of these items in the range of 5-100 µg/mL. Neither the DMSO-BAR, HA-BAR, nor berberine had any effect at tested concentrations on bacterial growth or violacein production (Figure 1). All the concentrations were deemed suitable for further *in vivo* assays, as they all were non-inhibitory with respect to growth, and hence suitable for studying any possible effect on bacterial virulence.

**Figure 1.**
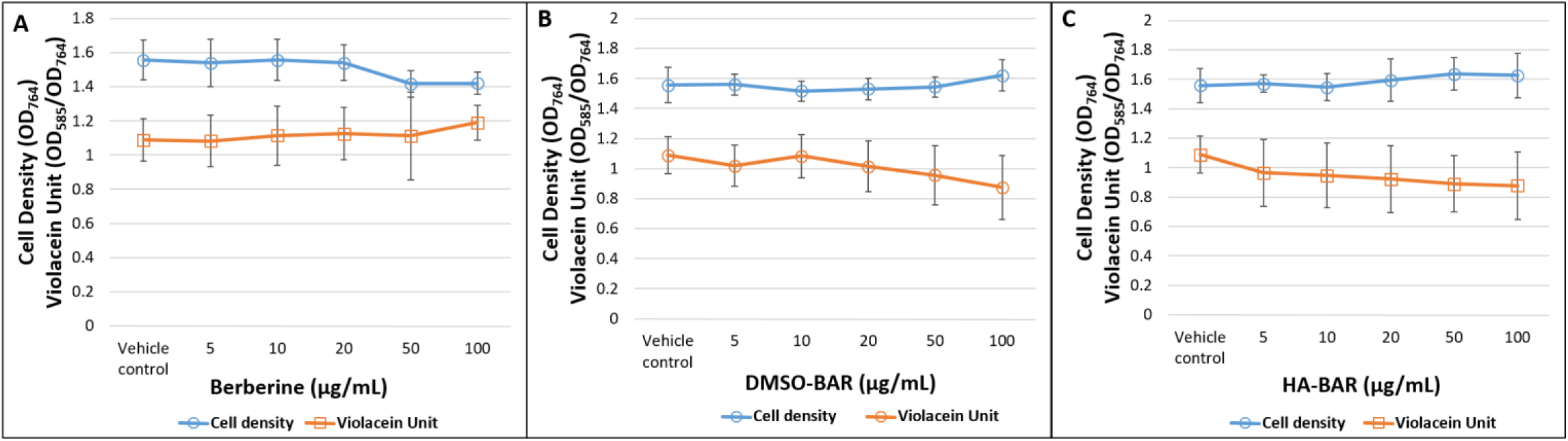
Neither berberine nor *B. aristata* affected growth and pigment production in *C. violaceum*. (A) Berberine; (B) DMSO-BAR, (C) HA-BAR OD of violacein was measured at 585 nm, and Violacein Unit was calculated as the ratio OD_585_/OD_764_ (an indication of violacein production per unit of growth); DMSO-BAR: DMSO-soluble fraction of *B. aristata* roots; HA-BAR: Hydroalcoholic extract of *B. aristata* roots; Control refers to the vehicle control containing 0.5%v/v DMSO, wherein DMSO did not affect growth and pigment production of *C. violaceum*. Ciprofloxacin (5 μg/mL) used as positive control inhibited bacterial growth completely.

### 3.2. *In vivo* experiments

#### 3.2.1. Anti-pathogenic assay

When the model host *C. elegans* was challenged with *C. violaceum* in presence of berberine or the plant extracts, the pathogen could kill lesser worms than in their absence (Figure-2; Supplementary videos: A-E). Since the concentrations of the extract or berberine tried were all growth non-inhibitory, the observed effect can be said to be growth-independent. Berberine and the plant extracts not only saved majority of the worms from pathogen-induced death, they also allowed worm reproduction. Ciprofloxacin, an antibiotic employed as positive control, though could save worms from pathogen-induced death, worms in antibiotic wells were not able to reproduce. In this respect, plant products can be said to have exerted better protective effect than ciprofloxacin. Berberine’s dual beneficial effect on worm survival as well as healthspan, in face of pathogen challenge is also supported by the higher worm activity in wells corresponding to berberine (Figure 3A). Berberine’s anti-pathogenic activity assumes even higher importance in light of the fact that of the six antibiotics belonging to three different classes, tried at same concentration as that of berberine (10 µg/mL), only ciprofloxacin was found to be active, and that too supported survival only, not fertility or active motility (Figure 3B). It is interesting to note that these all six antibiotics were selected based on their growth-inhibitory activity observed in the disc diffusion assay (Table S1).

**Figure 2.**
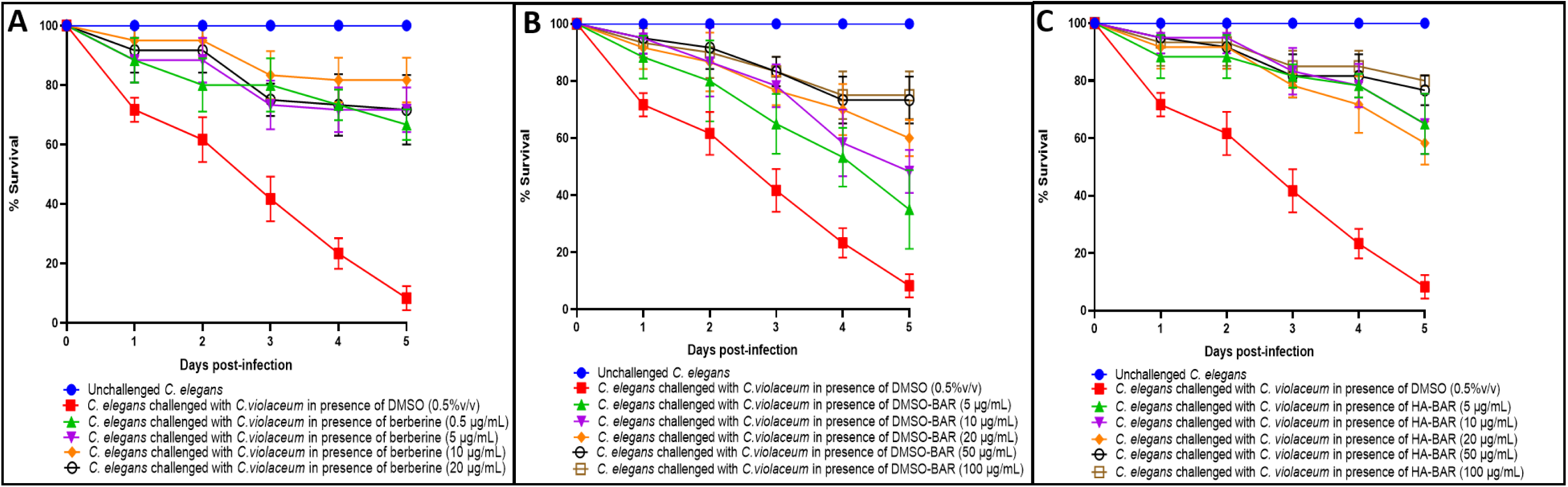
*C. violaceum*’s virulence towards the host worm gets attenuated in presence of berberine or *B. aristata* extracts. **(A) Berberine.** In face of pathogen-challenge, worm population registered 58.33%*** ± 5.16, 63.33%*** ± 7.5, 73.33%*** ± 7.5 and 63.33%*** ± 10.6 higher survival in presence of berberine at 0.5 μg/mL, 5 μg/mL, 10 μg/mL, and 20 μg/mL respectively. See supplementary videos A-E. **(B) DMSO-BAR.** In face of pathogen-challenge, worm population registered 26.66%*** ± 13.8, 40%*** ± 7.5 51.6%*** ± 6.3, 65%*** ± 8.16, and 66%*** ± 8.3 higher survival in presence of DMSO-BAR at 5 μg/mL, 10 μg/mL, 20 μg/mL, 50 μg/mL, and 100 μg/mL respectively. **(C) HA-BAR.** In face of pathogen-challenge, worm population registered 56.66%*** ± 10.4, 56.66%*** ± 10.4, 50%*** ± 7.5, 68.3%*** ± 5.16, and 70%*** ± 4 higher survival in presence of HA-BAR at 5 μg/mL, 10 μg/mL, 20 μg/mL, 50 μg/mL, and 100 μg/mL respectively. Berberine was tested at four different concentrations, and the both the extracts were tested at five different concentrations. Survival curve for berberine contain lines pertaining only to those concentrations, whose effect was statistically different from other concentrations tried. Of the five antibiotics (ampicillin, gentamicin, vancomycin, streptomycin, and ticarcillin; all at 10 µg/mL) tried as positive control, only ciprofloxacin was effective, which allowed 66.66%*** ± 5.77% higher worm survival in face of pathogen challenge. *** p ≤ 0.001

**Figure 3.**
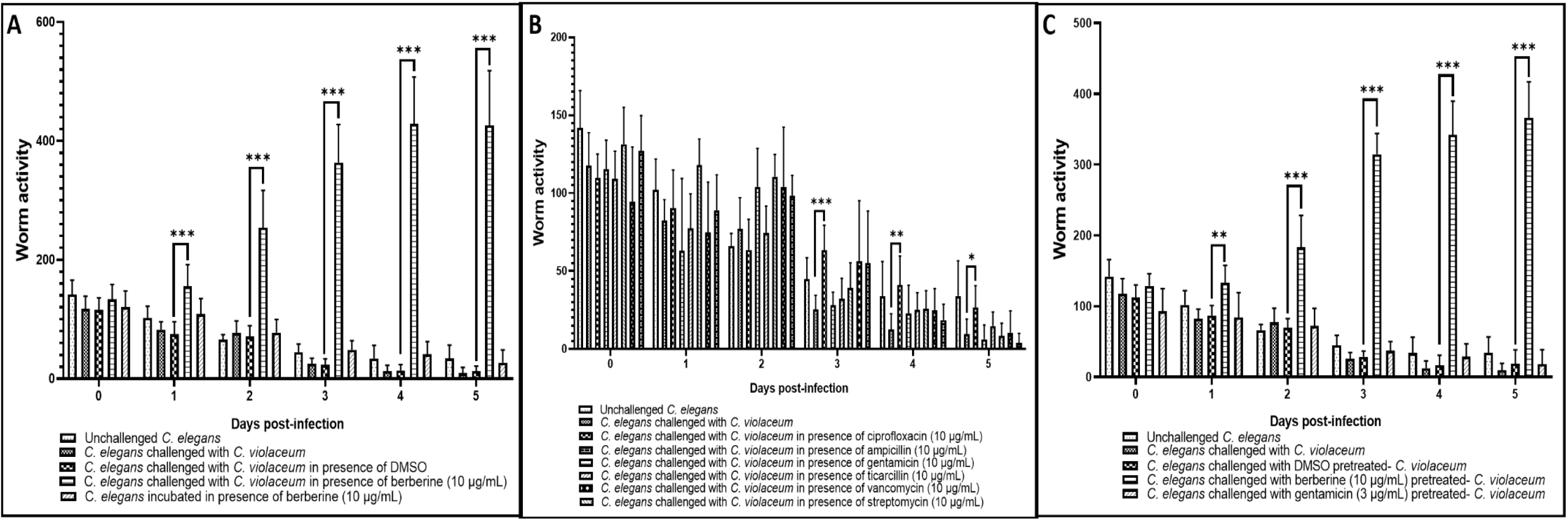
Berberine not only confers survival benefit on worms, but also supports better healthspan in face of pathogen challenge. Till second day, higher activity counts in wells pertaining to berberine indicates better worm motility, but not better survival, as death in worm population was observed only third day onward. However, notable increase in motility in berberine-wells on day three onward is also due to presence of progenies. Beneficial effect of berberine is observed in anti-pathogenic (A) as well as anti-virulence (C) assay. Failure of gentamicin in protecting worms from *C. violaceum* is also visible here. (B) Ciprofloxacin was the most effective antibiotic against *C. violaceum*. To make a direct comparison, all antibiotics and berberine were tested at the same concentration (10 µg/mL). However, except ciprofloxacin, no other antibiotic could rescue worms from pathogen-mediated death. Worm activity plotted here, a surrogate for their survival as well as healthspan, was quantified through an automated worm tracker (WMicroracker^®^; Phylumtech ARENA). Assays were conducted in 24-well microplate format. Temperature of the plate-holding chamber of worm tracker was set at 23°C. Data acquisition was programmed at 15 min intervals, allowing time-lapse tracking of worm movement and responses. Values reported here are means of two independent experiments, each containing three replicates. *p<0.05; **p<0.01; ***p<0.001

While all the three plant products were able to attenuate virulence of the pathogen, berberine at lower concentrations was marginally superior than DMSO-BAR or HA-BAR. At 10 µg/mL, berberine was at par to ciprofloxacin, as per fifth day end-point. At the lowest concentration (5 µg/mL) tried, DMSO-BAR was lesser effective than HA-BAR as well as berberine. Latter showed anti-pathogenic efficacy at a concentration as low as 0.5 µg/mL.

Since in this assay, both the host as well as the pathogen were simultaneously exposed to the test extracts/compound, we further performed an anti-infective assay (results described in next section) to investigate whether the better worm survival observed in the anti-pathogenic assay was due to plant product’s effect on the pathogen alone.

#### 3.2.2 Anti-infective assay

When *C. violaceum* was pre-grown in berberine- or extract-supplemented media before being allowed to attack the host worms, its virulence was found to be attenuated (Figure 4). Based on fifth-day end-point, berberine’s virulence-attenuating effect was not dose-dependent (Figure 4A; Supplementary videos: F-H), as 10 µg/mL of it was found to attenuate bacterial virulence more than the higher concentrations. Maximum anti-virulence effect of DMSO-BAR and HA-BAR was observed at 50 µg/mL and 100 µg/mL respectively. Berberine’s anti-virulence activity was also evident from the higher activity recorded in worm population challenged with berberine-pre-treated pathogen (Figure 3C).

**Figure 4.**
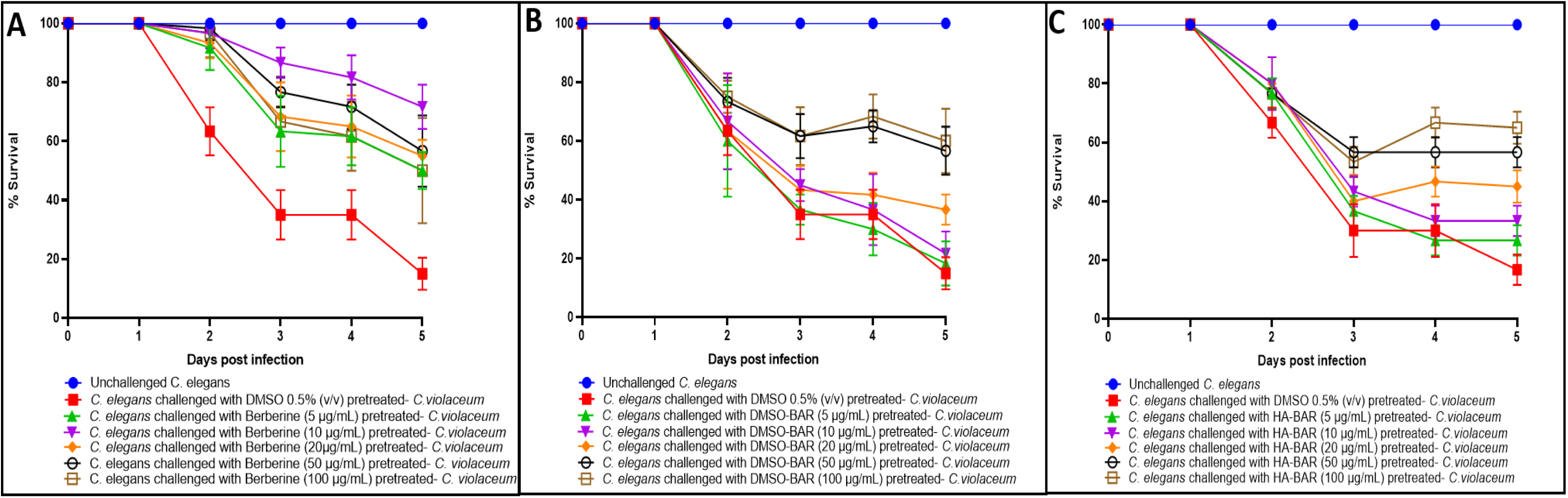
*C. violaceum* grown in media supplemented with berberine or *B. aristata* extract displays attenuated virulence towards the model host *C. elegans*. Berberine (A), DMSO-BAR (B), or HA-BAR (C) pre-treatment reduced bacterial virulence towards *C. elegans.* (A) Pre-treatment of *C. violaceum* with berberine at 5,10,20, 50 and 100 μg/mL reduced its virulence towards the host worm by 50%*** ± 6.32, 71.60%*** ± 7.52, 55%*** ± 5.47, 56.6%*** ± 12.10, and 50%*** ± 17.8 respectively. See supplementary videos F-H. **(B)** Pre-treatment of *C. violaceum* with DMSO-BAR at 20, 50 and 100 μg/mL reduced its virulence towards the host worm by 36.6%*** ± 5.16, 56.6%*** ± 8.16, and 60%*** ± 10.9 respectively. **(C)** Pre-treatment of *C. violaceum* with HA-BAR at 5, 10, 20, 50 and 100 μg/mL reduced its virulence towards the host worm by 26.6%*** ± 5.16, 33.3%*** ± 5.16, 45%*** ± 5.47, 56.6%*** ± 5.16, and 65%*** ± 5.16 respectively. These percent values refer to the difference between number of worms surviving in experimental and control wells. Progenies were observed on third day in all those experimental wells, wherein reduction in bacterial virulence was observed. Gentamicin (3 μg/mL) and vancomycin (7 μg/mL) tried as positive control at IC_50_ could not reduce bacterial virulence. ***p<0.001

Putting together the results of anti-infective and anti-pathogenic assays, it can be said that ability of all the three test items to protect worms from *C. violaceum* stems largely from their virulence-attenuating effect on the pathogen, and any immunomodulatory effect on the host seems not to be involved. Further confirmation of this interpretation in case of berberine was obtained through a prophylactic assay too (Figure S2). Since in both these assays, berberine was found to be better than both DMSO-BAR and HA-BAR, further assays were performed with berberine only.

#### 3.2.3. Post-infection assay

To investigate whether berberine can rescue already-infected worms, we first allowed *C. violaceum* to establish infection for 24 h or 48 h in the worm population, and then added berberine into the assay system as a possible post-infection therapeutic. Berberine was able to rescue worms when added within 24 h post-infection, but not when added later than that. (Figure 5; Supplementary videos: I-J).

**Figure 5.**
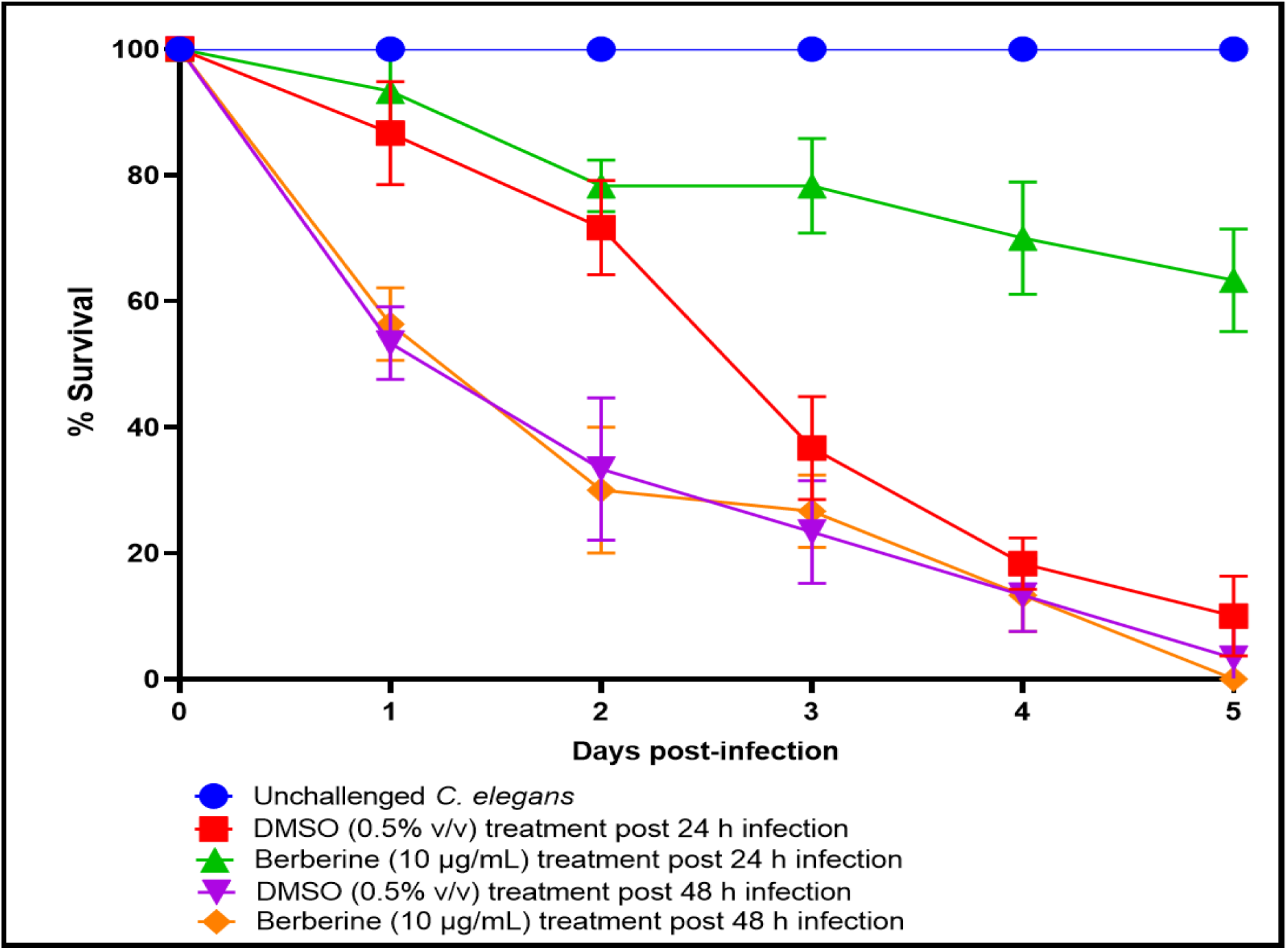
Berberine can rescue worms from pathogen challenge if added within 24 hour post-infection. While DMSO (0.5%v/v) had no effect on post-infection survival of worms, berberine at 10 µg/mL could rescue 53.33%***±8.16 worms (fifth day endpoint) if added within 24 hours of pathogen challenge. Late addition of berberine (48 h post-infection) had no protective effect on worms. Ciprofloxacin (10 µg/mL) tried as a positive control could rescue worms up to a small extant, if added within 18 hours of pathogen challenge. However, its protective effect was 46% lesser (p=0.003) than that of berberine, and was sustained only till fourth day. Line corresponding to ciprofloxacin is not included in this graph to avoid overcrowding. ***p<0.001

Once berberine’s ability to attenuate *C. violaceum*’s virulence was confirmed, we tried to get some insight into empirical mode of action of worm killing by this pathogen, and where berberine interferes. When we challenged worms with *C. violaceum* cells alone or culture supernatant alone, it was found that cell-mediated killing is the major mechanism in this host-pathogen interaction. When these assays were repeated in presence of berberine, this phytocompound was able to save worms from cells-only challenge as well as whole bacterial culture challenge (Figure 6; Supplementary videos: K-M).

**Figure 6.**
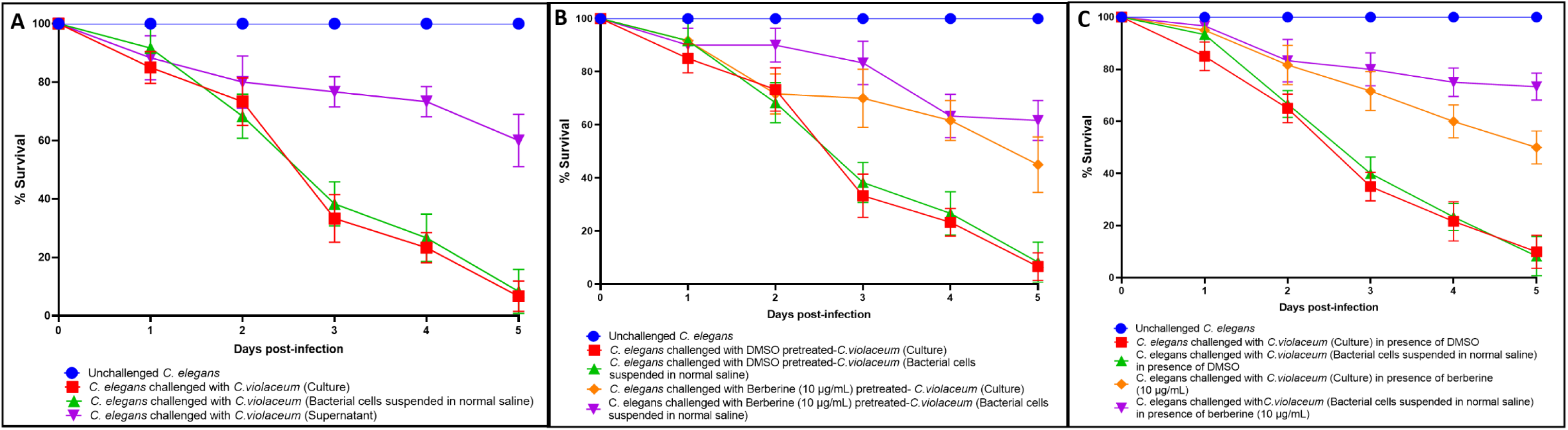
Berberine protects *C. elegans* against the cell-mediated toxicity exerted by *C. violaceum*. (A) Magnitude of cell-mediated toxicity exerted by *C. violaceum* against worms is much higher that supernatant-mediated toxicity. Worms in presence of bacterial culture supernatant are able to reproduce, and some of them died when progenies exited the parent body. Parallel running of the lines corresponding to whole culture and cells-only indicates that cell-mediated killing is the major mode of *C. violaceum*’s virulence towards the worms. Berberine’s protective effect on worms, against the cell-mediated toxicity, was demonstrated in anti-virulence (B) as well as anti-pathogenic (C) assay. See supplementary videos K-M.

Berberine was found to be more effective against *C. violaceum*, when worms are challenged with the pathogen in presence of berberine during the anti-pathogenic assay, as compared to its efficacy in the anti-infective or post-infection assay. After confirming berberine’s anti-virulence activity against *C. violaceum*, we performed two more *in vivo* assays to answer the questions: Whether berberine exerts any post-exposure effect on this pathogen? Whether *C. violaceum* can develop resistance against berberine after repeated exposure to this phytochemical? While no post-exposure effect was observed (Figure S3), results of the assay pertaining to the latter question are described in the next section.

### 3.2.4. Repeated exposure of *C. violaceum* to berberine did not induce resistance

Since development of resistance by the target pathogen against antibiotics is a major reason limiting their utility, to check whether *C. violaceum* can develop resistance to berberine’s virulence-attenuating activity, we subcultured *C. violaceum* in berberine (10 or 20 µg/mL)-supplemented medium. After ten such subculturing events, the ‘berberine-habituated’ *C. violaceum* was still not able to exert virulence at par to its berberine-not-exposed counterpart (Figure 7). *C. violaceum* was grown for multiple generations in presence of berberine, and despite the presence of this selection pressure, the pathogen could not develop resistance to berberine’s anti-virulence activity. One of the reasons for this may be the fact that berberine at the concentrations tried here (10-20 µg/mL) has no inhibitory effect on bacterial growth, and the magnitude of ‘selection pressure’ exerted by a growth non-inhibitory agent may not be sufficient to trigger resistance development.

**Figure 7.**
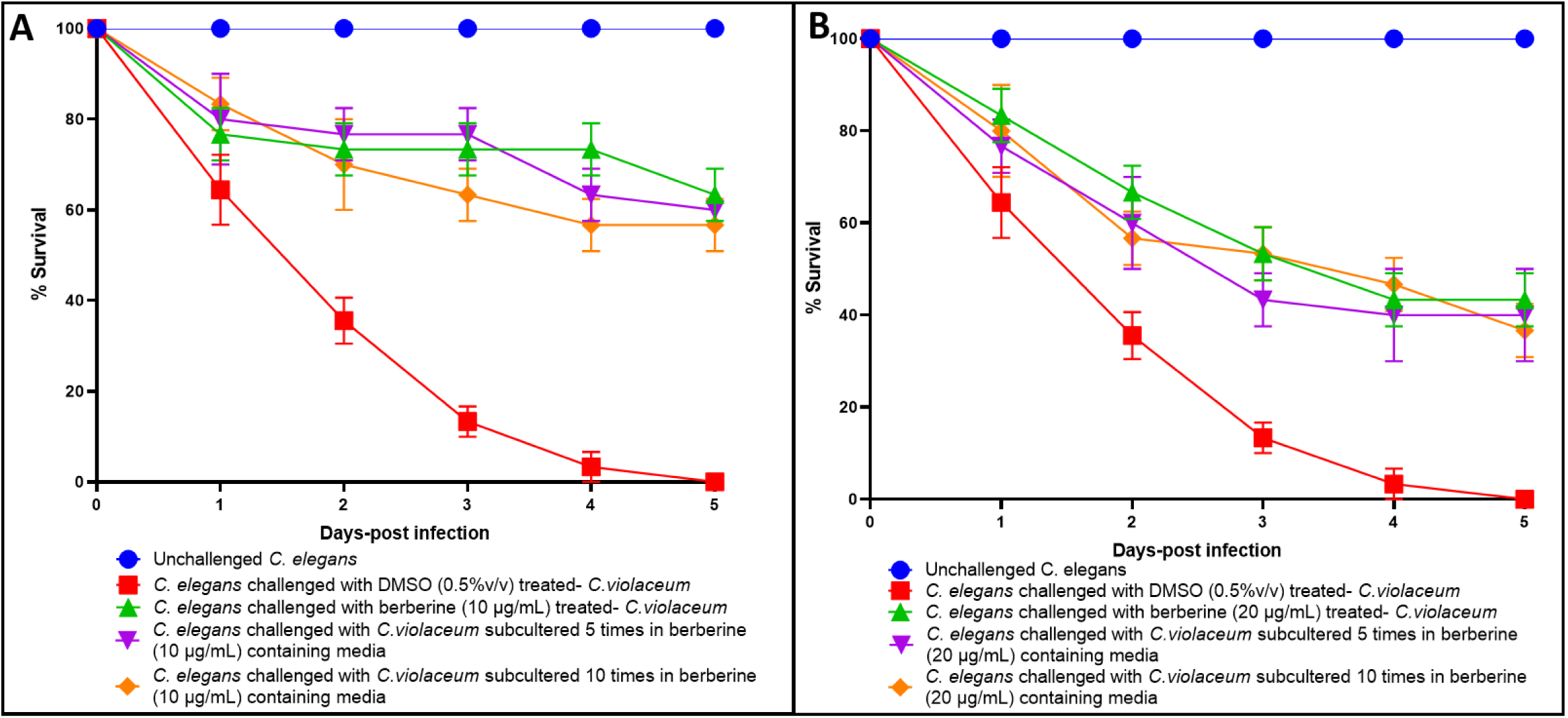
*C. violaceum* did not develop resistance even after repeated exposure to berberine. **(A)** *C. violaceum* obtained after five or ten subculturing in berberine (10 μg/mL)-containing media experienced 60%***±0 and 56%**±5.77 reduction in virulence respectively. (B) *C. violaceum* obtained after five or ten subculturing in berberine (20 μg/mL)-containing media experienced 40%***±10 and 36.66 %***±5.77 reduction in virulence respectively. None of the values for berberine-habituated cultures were statistically different than those for control culture (having single pre-exposure to berberine). ***p<0.001

### 3.3. *In vitro* results

After confirming berberine’s anti-virulence activity at non-growth inhibitory concentration, its effect on different virulence traits of *C. violaceum* was assessed through appropriate *in vitro* assays.

#### 3.3.1. Berberine compromises haemolytic potential of *C. violaceum*

Supernatant from the berberine-exposed *C. violaceum* culture was less haemolytic than that obtained from the control culture. Berberine at 10-100 µg/mL was able to compromise the ability of this pathogen’s culture supernatant to lyse human red blood cells by 22-47% (Figure 8).

**Figure 8.**
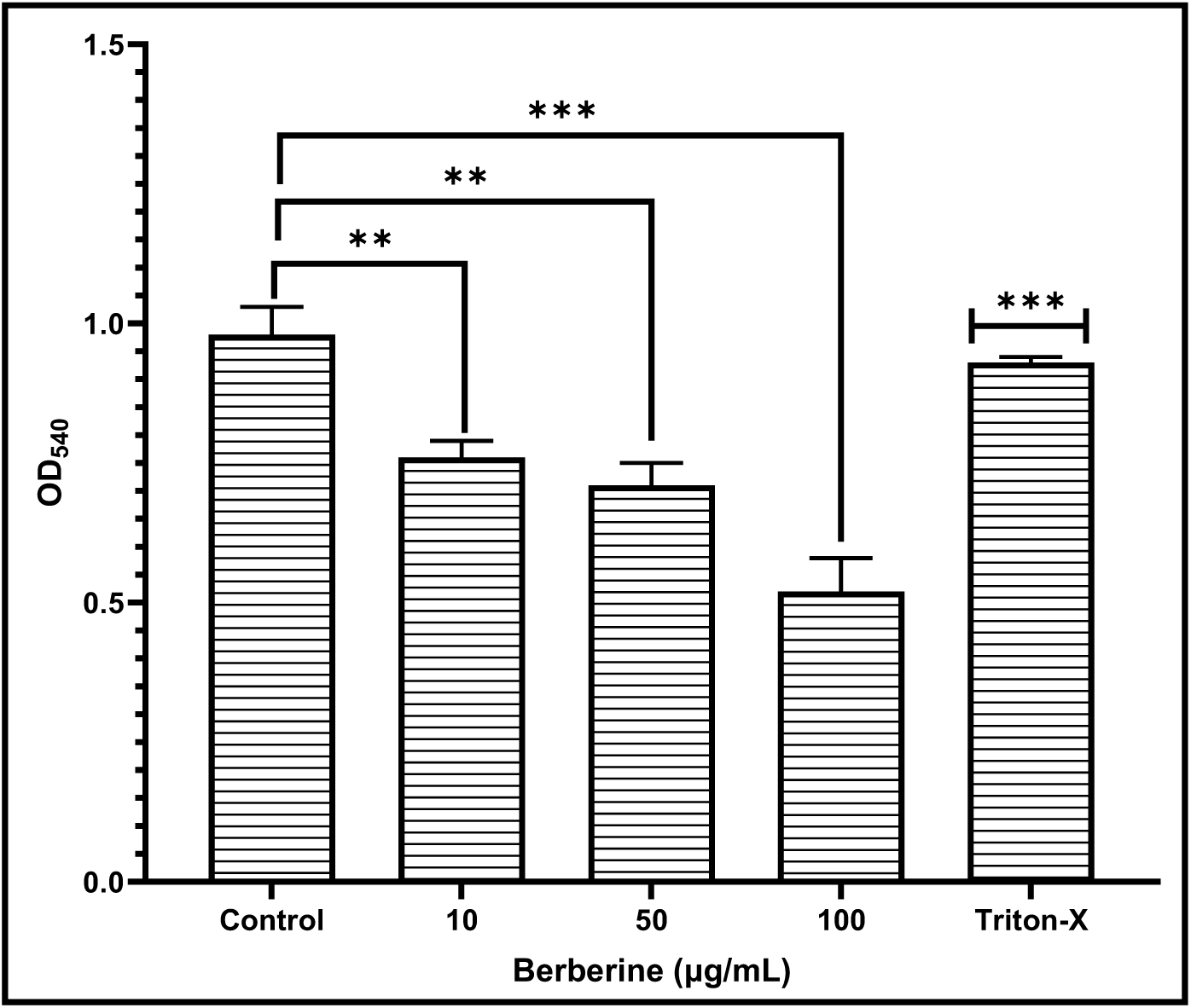
Berberine curbed haemolytic potential of *C. violaceum*. Hemoglobin released as a result of haemolysis was quantified as OD_540_; 1% triton and PBS (pH 7.4) were used as positive and negative control respectively; **p<0.01; ***p<0.001

#### 3.3.2. Berberine forces higher exopolysaccharide (EPS) production in *C. violaceum*

*C. violaceum*, when grown in presence of different concentrations of berberine, produced more (31-43%↑) EPS per unit of cell density (Figure 9A). Since EPS is the major ingredient of biofilm matrix, this observation corroborates well with result of the biofilm assay (Figure 9B), wherein biofilm biomass was found to be higher in case of *C. violaceum* grown in presence of berberine. Higher biofilm biomass/enhanced EPS production is considered as an indication of the stress in bacterial population (Mohammed et al., 2020; Wang et al., 2020).

**Figure 9.**
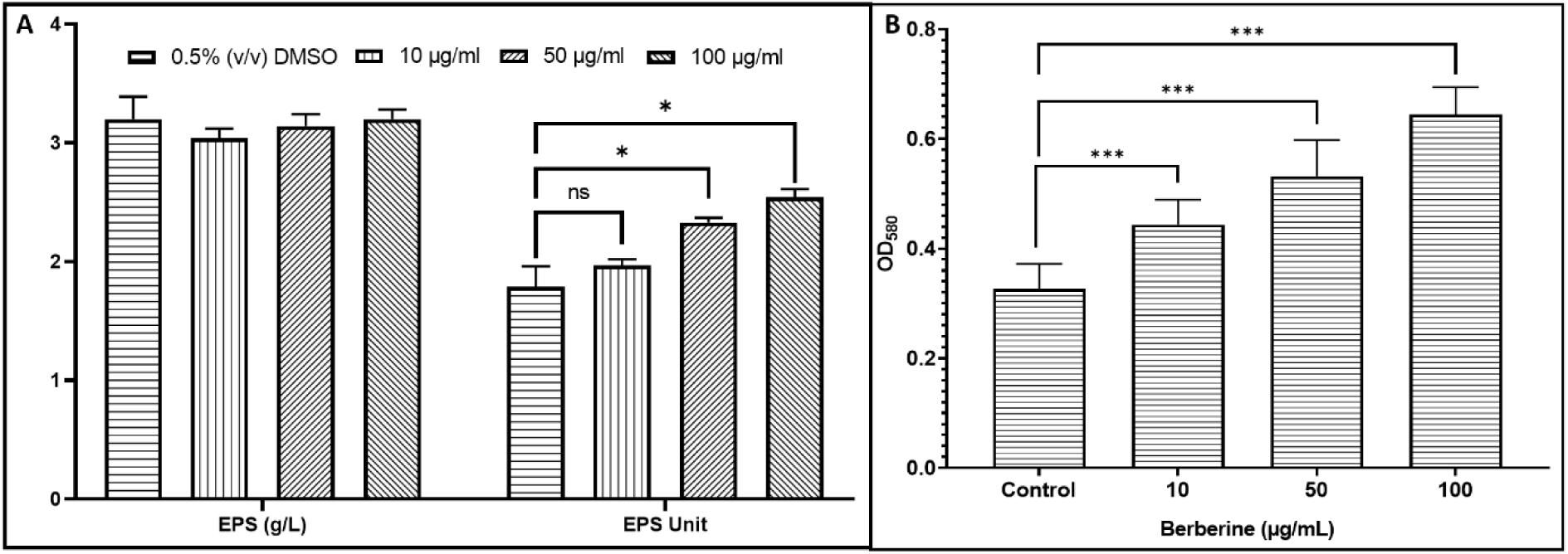
Berberine enhances exopolysaccharide (EPS) synthesis (A) and biofilm biomass (B) in *C. violaceum*. (A) EPS Unit was calculated as the cell density (OD_764_): EPS (g/L) ratio. (B) Biofilm biomass was quantified by measuring the crystal violet (OD_580_) trapped by the biofilm. *p<0.05; ***p<0.001; ns: not significant

#### 3.3.3. Berberine exerts an inhibitory effect on efflux machinery of *C. violaceum*

When *C. violaceum* was challenged with EtBr in presence or absence of berberine, the bacterial cells accumulated more EtBr in presence of berberine than that in its absence (Figure 10). EtBr being a DNA-damaging toxin, bacterial population responds to its presence by upregulating their efflux machinery, as a detoxification effort. Since bacterial cells accumulated more EtBr in presence of berberine, this phytocompound can be said to be acting as an efflux inhibitor. However, its efflux-inhibitory activity was observed at lower concentrations (10-50 µg/mL), and not at higher concentration (100 µg/mL). Berberine could exert efflux-inhibitory effect at par to the known efflux inhibitor, reserpine (used as a positive control), at half of the concentration used for the latter. This observation corroborates with the differential regulation (discussed later) of *norM* (coding for a MATE family efflux transporter; Table-2) in *C. violaceum* grown in presence of berberine. Berberine’s efflux-inhibitory effect seems to have been responded by bacterial effort to upregulate efflux-associated genes. Efflux inhibition is being viewed as a potent anti-virulence strategy owing to vital role of the efflux pumps in detoxification. Disturbance of efflux machinery positively or negatively, both is likely to prove detrimental to bacterial fitness, given the important functional role of efflux pumps in bacterial physiology (Sun et al., 2014). Since efflux pumps are involved in a broad range of physiological functions, and their expression is tightly regulated, efflux-inhibitors are looked as a promising intervention for treatment of bacterial infections.

**Figure 10.**
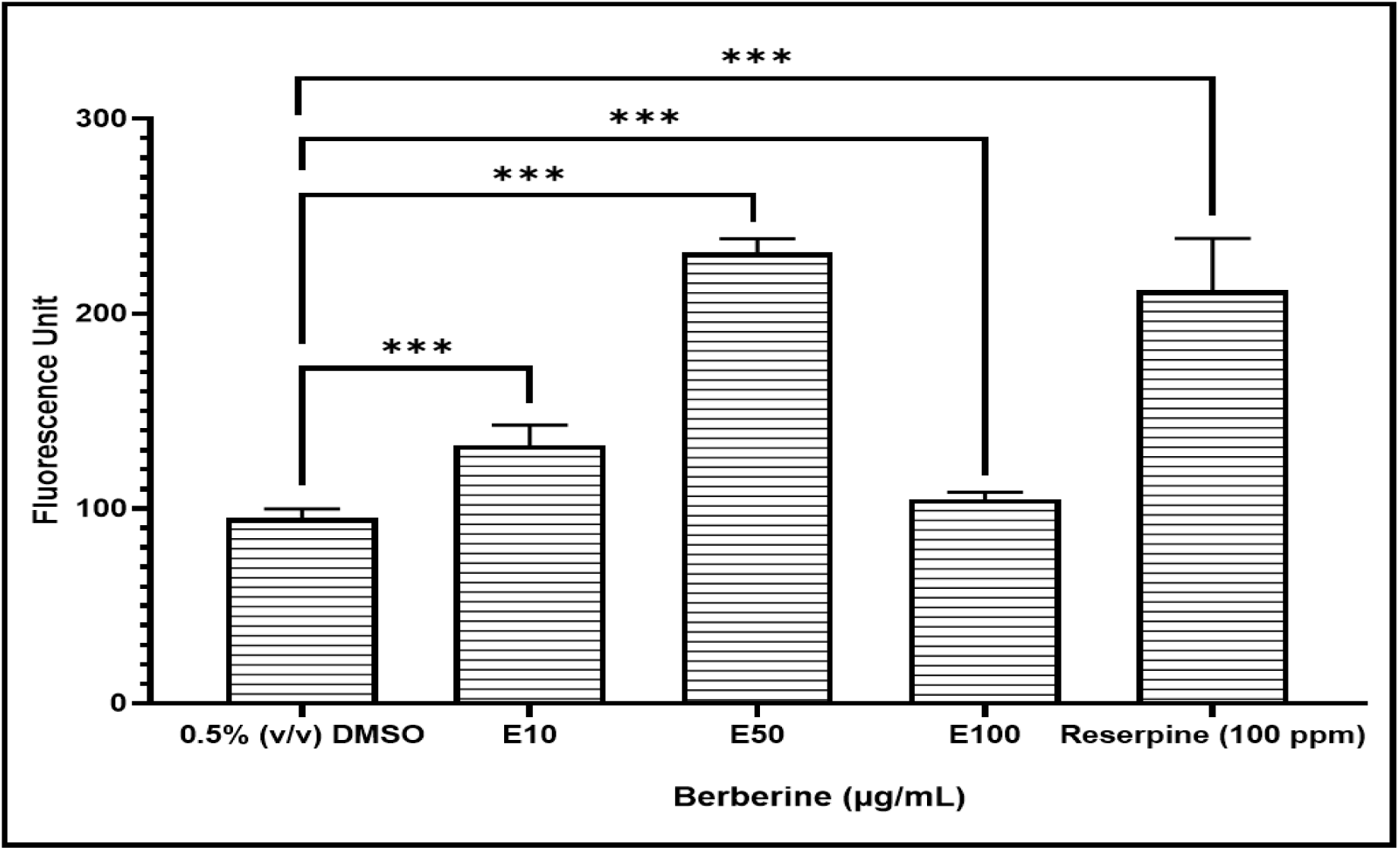
Berberine triggers differential regulation of efflux machinery in *C. violaceum*. Fluorescence of the intracellularly accumulated EtBr was 39%*** ± 10.3 and 142.9%*** ± 7 higher in *C. violaceum* exposed to berberine (10 µg/mL and 50 µg/mL, respectively). However, berberine’s efflux-inhibitory effect disappeared at higher concentration (100 µg/mL). Higher (122.4%*** ± 26.6) intracellular accumulation of EtBr, as expected, was observed in *C. violaceum* exposed to a known efflux inhibitor, reserpine, used as positive control. ***p<0.001; ns: not significant

### 3.4. Differential gene expression in berberine-exposed *C. violaceum*

To gain mechanistic insights into berberine’s anti-virulence activity against *C. violaceum*, we compared the gene expression profile of *C. violaceum* grown in presence of berberine with that grown in its absence, at the whole transcriptome level. We found a total of twelve genes (Table-2) expressed differently in berberine-exposed *C. violaceum* passing the dual cut-off of Log FC≥2 and FDR≤0.05. Of these twelve, 9 genes were upregulated and 3 were downregulated. The corresponding heat map (Figure S4) volcano plot (Figure S5), and MA plot (Figure S6) are provided in the Supplementary Materials.

**Table 2.** List of DEG in berberine-exposed C. violaceum satisfying the dual criteria of log fold-change ≥2 and FDR≤0.05.

| Sr. No | Feature ID | Gene | Product/function | Up-/Down-regulation | Log FC |
| --- | --- | --- | --- | --- | --- |
| 1 | CV_RS12005 | *norM* | MATE family efflux transporter | ↑ | 5.70 |
| 2 | CV_RS17225 | *norB* | nitric-oxide reductase large subunit | ↓ | 4.88 |
| 3 | CV_RS05670 | *zntA* | heavy metal translocating P-type ATPase | ↑ | 4.81 |
| 4 | CV_RS12000 | *pepQ* | Xaa-Pro peptidase family protein | ↑ | 4.72 |
| 5 | CV_RS12010 | *tetR* | TetR family transcriptional regulator | ↑ | 3.39 |
| 6 | CV_RS05695 | A0A202BDG5 | GtrA family protein | ↑ | 3.22 |
| 7 | CV_RS22050 | NA | putative metalloprotease CJM1_0395 family protein | ↑ | 3.20 |
| 8 | CV_RS09795 | *nirK/aniA* | copper-containing nitrite reductase | ↓ | 3.13 |
| 9 | CV_RS05680 | A0A202BDM3 | acyl-CoA thioesterase | ↑ | 2.90 |
| 10 | CV_RS12015 | Q7NV87 | ABC transporter permease | ↑ | 2.69 |
| 11 | CV_RS01270 | A0A1R0MRG2 | DUF3079 domain-containing protein | ↓ | 2.69 |
| 12 | CV_RS05700 | A0A202BDS7 | glycosyltransferase family 2 protein | ↑ | 2.67 |

Genes are arranged in decreasing order of log fold (Log FC) change. Databases consulted for gene functions were; KEGG (Kyoto Encyclopedia of Genes and Genomes: https://www.genome.jp/kegg/); Uniprot (https://www.uniprot.org/). NA: not available

Among all the differently expressed genes (DEG), *norM* coding for MATE (Multidrug And Toxic Compound Extrusion) family efflux transporter topped the list with respect to the magnitude (5.70↑) of differential expression. Three other genes, CV_0385, coding for a *tetR* family transcriptional regulator; one coding for heavy metal translocating P-type ATPase (*zntA*); and Q7NV87, coding for a ABC transporter permease, associated with detoxification/efflux/secretion/transport were also upregulated. This taken together with the experimentally observed efflux-inhibitory effect of berberine (Figure-10), indicates that the pathogen has tried to activate its stress-response machinery to encounter the efflux-inhibitory/toxic effect of berberine. All these four genes have been known to be associated with expression of MDR (multidrug resistance) efflux pumps in different pathogenic bacteria (Grangeiro et al., 2004; Kumar et al., 2013; Thornhill, 2016), and hence though used at growth non-inhibitory concentration, berberine seems to have induced a response in *C. violaceum* similar to that induced by an antibiotic. *C. violaceum*’s response to berberine seems to have some similarity to its response to heavy-metal toxicity, as this bacterium is known to try for enhancing its biofilm biomass and efflux expression when challenged with heavy metals (Silver and Ji, 1994). Upregulation of *zntA*, which is believed to confer resistance to xenobiotics (Brazilian National Consortium, 2003) is an additional indication to believe that the pathogen is under stress in presence of berberine. Expression of the majority of the efflux pumps is subject to tight control by various transcriptional regulators, underlying their roles in facilitating the adaptation of bacteria to specific stimuli. Overexpression of efflux pump has been shown to impair the fitness and virulence of pathogens like *S. maltophilia* (Alonso et al., 2004; Andersson et al., 2007). Improper overexpression of efflux pumps may cause unwanted efflux of metabolites or other signaling molecules, resulting in deleterious effects on cell physiology (Sun et al., 2014).

Among the overexpressed efflux-associated genes, *norM* is involved in expulsion of a broad range of structurally diverse toxic substances like 4′,6-diamidino-2-phenylindole dihydrochloride (DAPI), ciprofloxacin, norfloxacin, acriflavine, kanamycin, streptomycin, daunomycin, doxorubicin, trimethoprim, and ethidium bromide (Huda et al., 2001). This efflux transporter or its homologues are also reported in other bacteria like *Vibrio parahaemolyticus*, *Escherichia coli* (Morita et al., 1998), and *Neisseria gonorrhoeae* (Long et al., 2008). Interference of efflux pumps like norM, which has poly-substrate specificity, may force the bacteria to expel some required metabolites too. Efflux pump activity may affect bacterial virulence indirectly, e.g. through altering quorum sensing responses, since bacterial MDR efflux pump do have role in autoinducer traffic. Hyperexpression of efflux system may result in lesser production of AHL autoinducers, causing the bacteria to produce lesser QS-controlled extracellular virulence factors. Active efflux of auto-inducers can limit their intracellular concentration and consequently its dependent activation of genes coding for virulence factors. The capability of efflux pumps to export QS autoinducers is supported by various studies, and in context of the present study, we may speculate that disturbed expression of the efflux machinery in berberine-challenged *C. violaceum* might have reduced the intracellular concentration of the autoinducers, not allowing effective regulation of virulence in a population density-dependent fashion.

Second most upregulated detoxification-related gene was *zntA*, which has been shown in other gram-negative bacteria like *Vibrio parahaemolyticus* to be crucial for maintenance of metal (Zn and Cd) homeostasis, oxidative stress resistance, and virulence (Zheng et al., 2024). Since *zntA* is already known to be induced by Zn, Cu, Co, Ni, and Cd, berberine can be said to induce stress in bacteria similar to that induced by toxic heavy metals. Improper functioning of *zntA* has also been shown in case of *Escherichia* coli to make it more susceptible to host-mediated killing (Barisch et al., 2018).

Another significantly upregulated efflux-associated gene was a *tetR* family transcriptional regulator. TetR family regulators are known to be involved in regulation of different MDR efflux pumps in various bacteria. Besides antibiotic resistance, the TetR regulators are widely associated with the regulation of genes encoding small-molecule exporters, metabolism, osmotic stress, homeostasis, biosynthesis of antibiotics, multidrug resistance, efflux pumps, enzymes implicated in different catabolic pathways, virulence and pathogenicity, modification and clearance of toxic compound, quorum sensing, and many other aspects of prokaryotic physiology (Cuthbertson and Nodwell, 2013). Seventeen proteins of TetR family are known in *C. violaceum*. Since TetR family typically comprises of transcriptional repressors (Ramos et al., 2005), its observed upregulation in berberine-challenged *C. violaceum* can be expected to repress expression of multiple traits regulated by this family. In this study, two such traits i.e. efflux (Figure-10), and haemolysis (Figure-8) were found to be compromised in *C. violaceum* in presence of berberine. In context of the latter, it is important to note that TetR proteins are already known to be involved in heme homeostasis (Hashimoto et al., 1997; Lechardeur et al., 2012). Berberine seems to have affected two of the most important functions regulated by the TetR family proteins, virulence and clearance of toxic compounds. Genes associated with these traits seems to have remained silent in presence of berberine, which corroborates with the attenuation of the berberine-pre-exposed pathogen’s ability to kill the model host *C. elegans* (Figure-4). Our results support the candidature of TetR family proteins as possible broad spectrum new drug targets. In particular, TetR-mediated multidrug efflux pump regulation is being viewed as potential target candidate in antibiotic-resistant bacteria. Since TetR is a chemical sensor used by bacteria to monitor cellular environmental dynamics, its disturbed expression can compromise the bacterial ability to monitor the level of intracellular toxic compounds (e.g. berberine in the present case). Given the fact that the intricate interplay between the transcriptional regulators and their cognate efflux pumps safeguard the intracellular homeostasis (Deng et al., 2013), berberine’s ability to alter the expression of transcription regulator as well as efflux machinery can be believed to disturb bacterial homeostasis.

The fourth trasnport-associated *C. violaceum* gene upregulated in presence of berberine was a probable ABC transporter permease (2.69 fold↑). While most of the ABC-type transport ORFs in C. violaceum are dedicated to nutrient acquisition, some are related to multidrug resistance. The wide spectrum of molecules/metabolites transported by ABC-type transport systems in C. violaceum also include nitrate/nitrite (Grangeiro et al., 2004), and this corroborates with disturbed expression of denitrification genes in our experimental culture.

Among other upregulated genes in the berberine-exposed *C. violaceum*, two (*pepQ* and a CJM1_0395 family protein) were coding for metalloproteases. Their dysregulation is important in light of the fact that a wide variety of pathological actions of bacterial metalloproteases are already documented in literature. In local bacterial infections, the metalloproteases may function as a decisive virulence determinant, while in systemic infections, they act as a synergistic virulence factor (Miyoshi and Shinoda, 1977). PepQ is a Xaa-Pro peptidase family protein. While not much is known about its function in *C. violaceum*, its association with antibiotic-resistance was demonstrated in *Mycobacterium tuberculosis*. Mutations in this gene were shown to confer low-level resistance to bedaquiline and clofazimine in *M. tuberculosis* (Almeida et al., 2016). Interaction of the metallopeptidase PepQ with secretion system proteins can influence functioning of the secretion system and bacterial virulence (Kuspa, 2014).

Upregulated genes also included a GtrA family protein. Though we could not find its specific role in *C. violaceum*, in other bacteria, members of the GtrA family are known to have role in functioning of secretion system (Xie et al.,2022) and carbohydrate metabolism (Kolly et al., 2015). One more upregulated gene, *tesA,* is a thioesterase required for outer-envelope lipid biosynthesis, and its validity as an effective pharmacological target has been demonstrated in mycobacteria (Nguyen et al., 2018). Its dysregulation can be thought to affect lipid and cell wall metabolism in the target pathogen, which in turn may influence its virulence. Despite the physiological functions of acyl-CoA thioesterases being an under-investigated area, they are believed to regulate lipid metabolism by maintaining appropriate concentrations of acyl-CoA, CoASH, and nonesterified fatty acids. In eukaryotes, their role is in the termination of fatty acid synthesis (Shahi et al., 2006). If they have the same function in *C. violaceum*, the observed upregulation of *tesA* might had a negative influence on fatty acid synthesis.

Last of the genes listed in Table-2 is an upregulated (2.67 fold ↑) glycosyltransferase family 2 protein. Glycosyltransferases are known to have important role in polysaccharide biosynthesis and biofilm formation (Becker et al., 2009). Upregulation of this genes corroborates with increased biofilm biomass formation in berberine-exposed *C. violaceum* observed in this study (Figure-9). Role of glycosyltransferases in bacterial fitness and pathogenicity has been demonstrated in multiple pathogens (Zhu et al., 2015; Ren et al., 2016).

Among the three downregulated genes, two (*nirK* and *norB*) belonged to the denitrification pathway. *nirK* codes for nitrite reductase, and its product nitric oxide (NO) is further converted into nitrous oxide by the nitric oxide reductase (nor). Since the magnitude of downregulation of *norB* (4.88↓) was more than that of *nirK* (3.13↓), it can be assumed that rate of NO production would have been more than rate of its clearance. NO is toxic in nature, and its build-up leads to nitrosative stress in the bacterial cell, which can have a negative impact on its overall fitness. Downregulation of the genes involved in denitrification pathway has earlier been shown by us in virulence-attenuated cultures of *Pseudomonas aeruginosa* (Joshi et al., 2019; Gajera et al., 2023; Parmar et al., 2024). Both the downregulated genes function downstream of nitrite, and hence the *C. violaceum* population in this case can be said to be experiencing nitrite accumulation in presence of berberine. Nitrite build-up can have negative impact on bacterial physiology, as it triggers the overexpression of efflux pumps and alters membrane permeability (Schreiber et al., 2006). Berberine’s growth-independent anti-virulence effect on *C. violaceum* observed in this study is consistent with Bazylinski et al. (1986)’s report that NO_2_^-^ reduction to N_2_O is not coupled to growth but may serve as a detoxification mechanism. Downregulation of the denitrification genes in the present case can be believed to have compromised the detoxification capacity of *C. violaceum*. Nitrate reduction in *Chromobacterium* species being coupled to energy conservation, downregulation of the denitrification pathway can have a negative impact on its overall fitness. Reduction of nitrite to nitrous oxide by *Chromobacterium* is viewed as an instance of nitrite detoxification, which is compromised in presence of berberine.

A protein-protein interaction (PPI) map of all DEG was generated (Figure-11), wherein only three genes (*norB, nirK*, and *zntA*) were found to have a non-zero node degree score. Since two of them belonged to the same pathway (i.e. denitrification), we chose to confirm their differential expression in berberine-challenged *C. violaceum* through PCR (Figure-12). A co-occurrence analysis (Figure 13) showed seven of the DEG identified in this study (including the two subjected to PCR-based validation) to have no homologue in humans. Exploring such targets further is logical, as antibacterial compounds targeting them are likely to exert selective toxicity towards the pathogen, without having adverse effect on humans.

**Figure 11.**
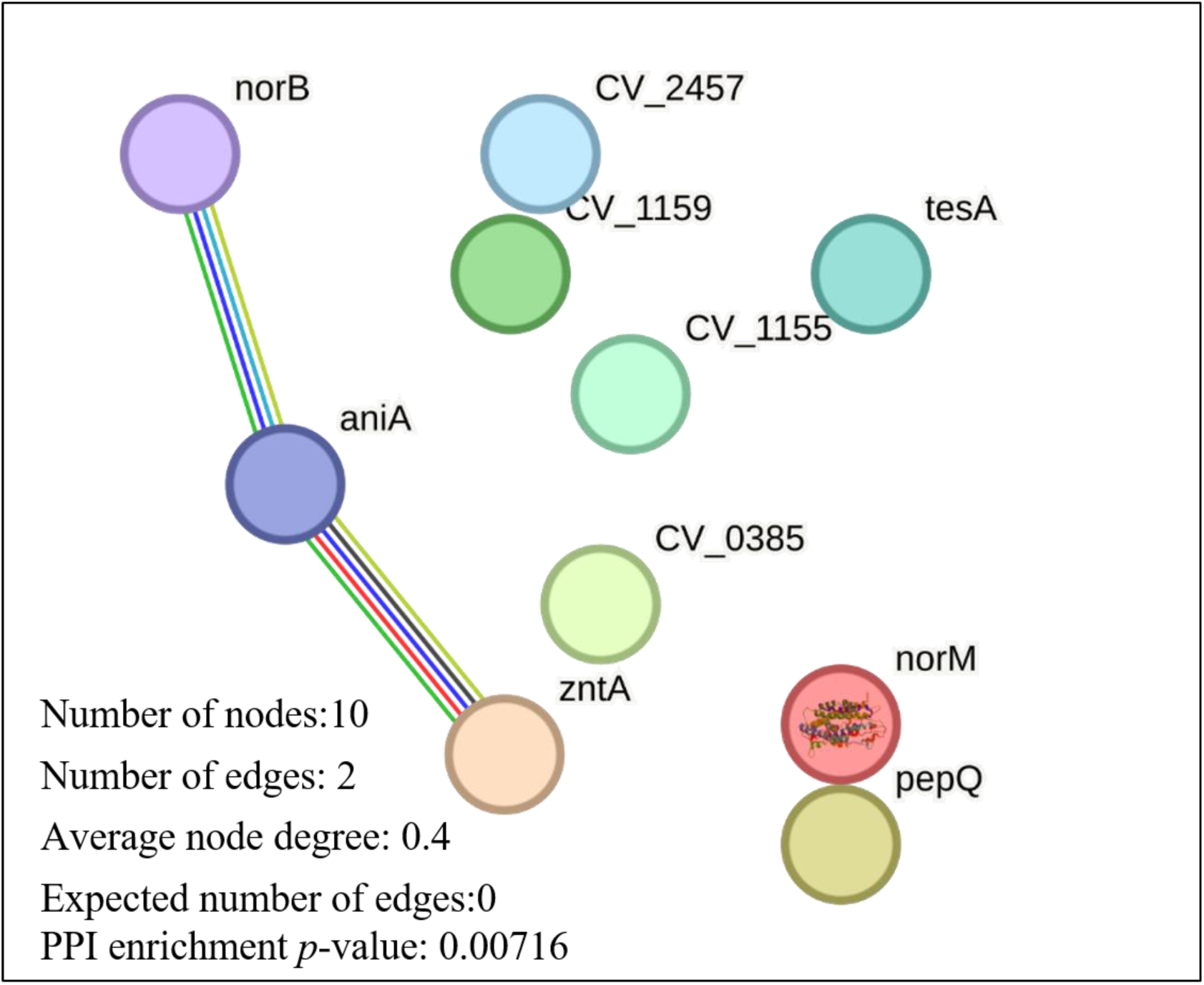
Protein-Protein Interaction (PPI) network of differentially expressed genes in berberine*-*treated *C. violaceum*. (https://string-db.org/cgi/network?taskId=bN4pFw9sGEZT&sessionId=blnLacFKaPLP)

**Figure 12.**
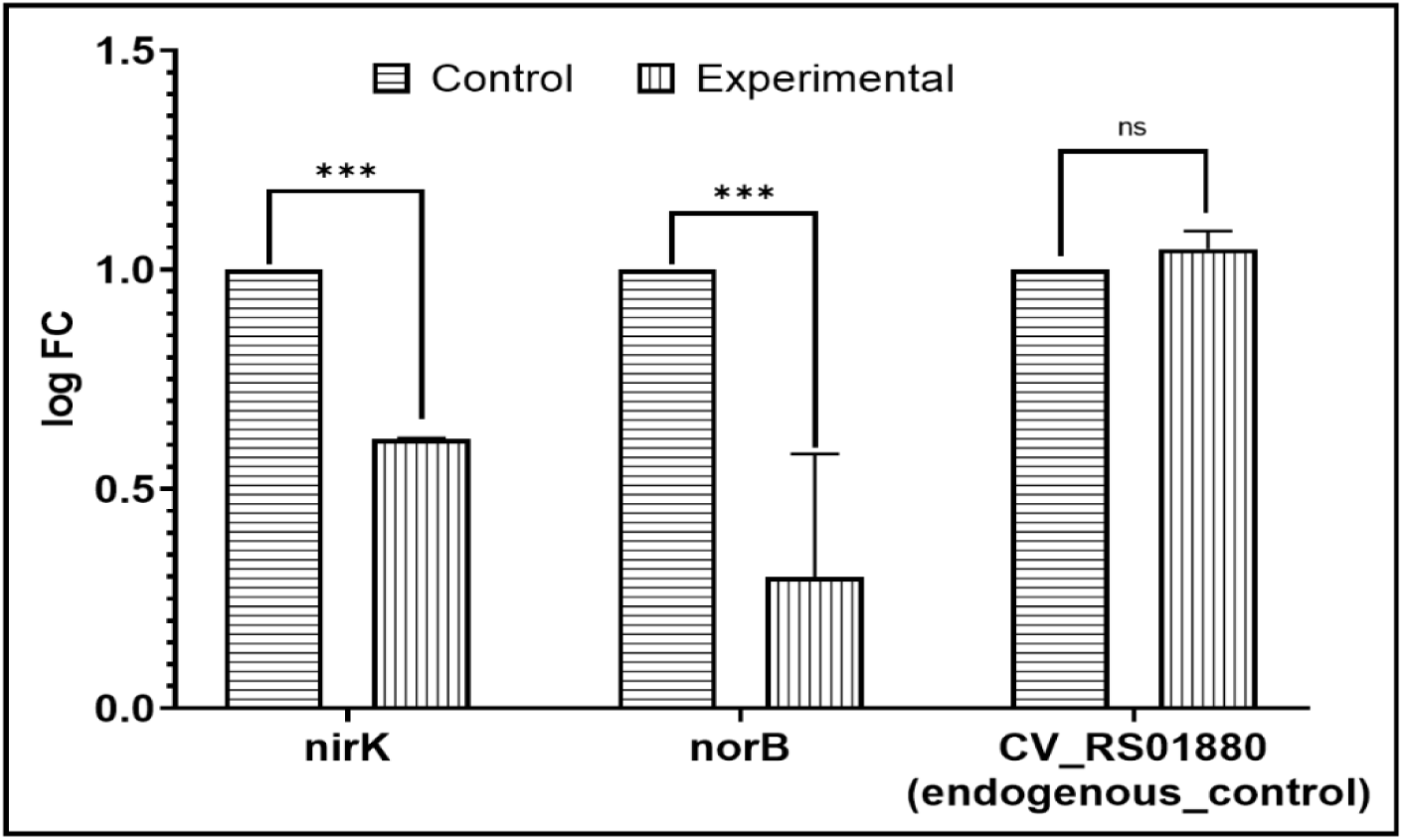
Confirmation of differential expression of selected downregulated genes in berberine-challenged *C. violaceum* using RT-PCR. *** p ≤ 0.001, ns: not significant; Corresponding gel images are provided in Supplementary Figure S7

**Figure 13.**
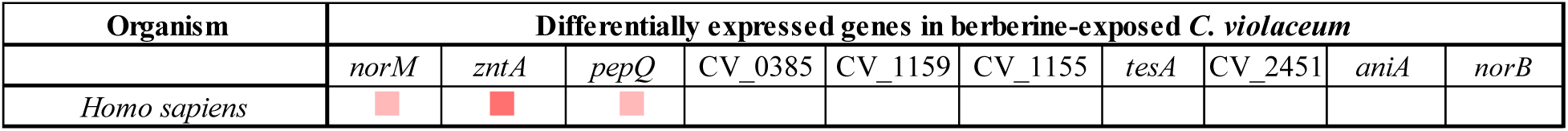
Co-occurrence analysis of genes coding for potential targets in *C. violaceum*, across *Homo sapiens* genome. The darker the shade of the squares, the higher the homology between the genes being compared.

In summary, root extract of *B. aristata* is capable of attenuating virulence of a multidrug resistant strain of *C. violaceum* against the model host *C. elegans*; and berberine appears to be the major active principle responsible for the observed anti-pathogenic activity. Berberine is able to influence multiple bacterial traits like virulence, efflux, EPS production and biofilm formation, and haemolysis by triggering differential expression of genes coding for efflux, denitrification, metalloproteases, etc. A schematic of various effects of berberine on *C. violaceum* is presented in Figure-14. Notably, berberine is able to reduce bacterial virulence by causing dysregulation of only a small number of genes, without exerting any growth-inhibitory effect, and perhaps that may be the reason why *C. violaceum* was not found to develop resistance to berberine (Figure-7), as the selection pressure exerted by berberine seems to be less. Results of the present study demonstrating anti-pathogenic activity of *B. aristata* root extract, and berberine at microgram levels validate the use of berberine in traditional medicine. For example, use of berberine-containing traditional Chinese medicine in treatment of diarrhea induced by bacterial infection, wherein berberine has been indicated to be safe for human use up to 100-300 mg/dose, thrice a day (Yu et al., 2020). Doses of berberine or *B. aristata* extracts in the range of 500–1500 mg/day are generally considered as safe by various regulatory agencies (Agarwal et al., 2025). Further investigation with respect to possible broad-spectrum anti-pathogenic activity of berberine against other important pathogens is warranted.

**Figure 14.**
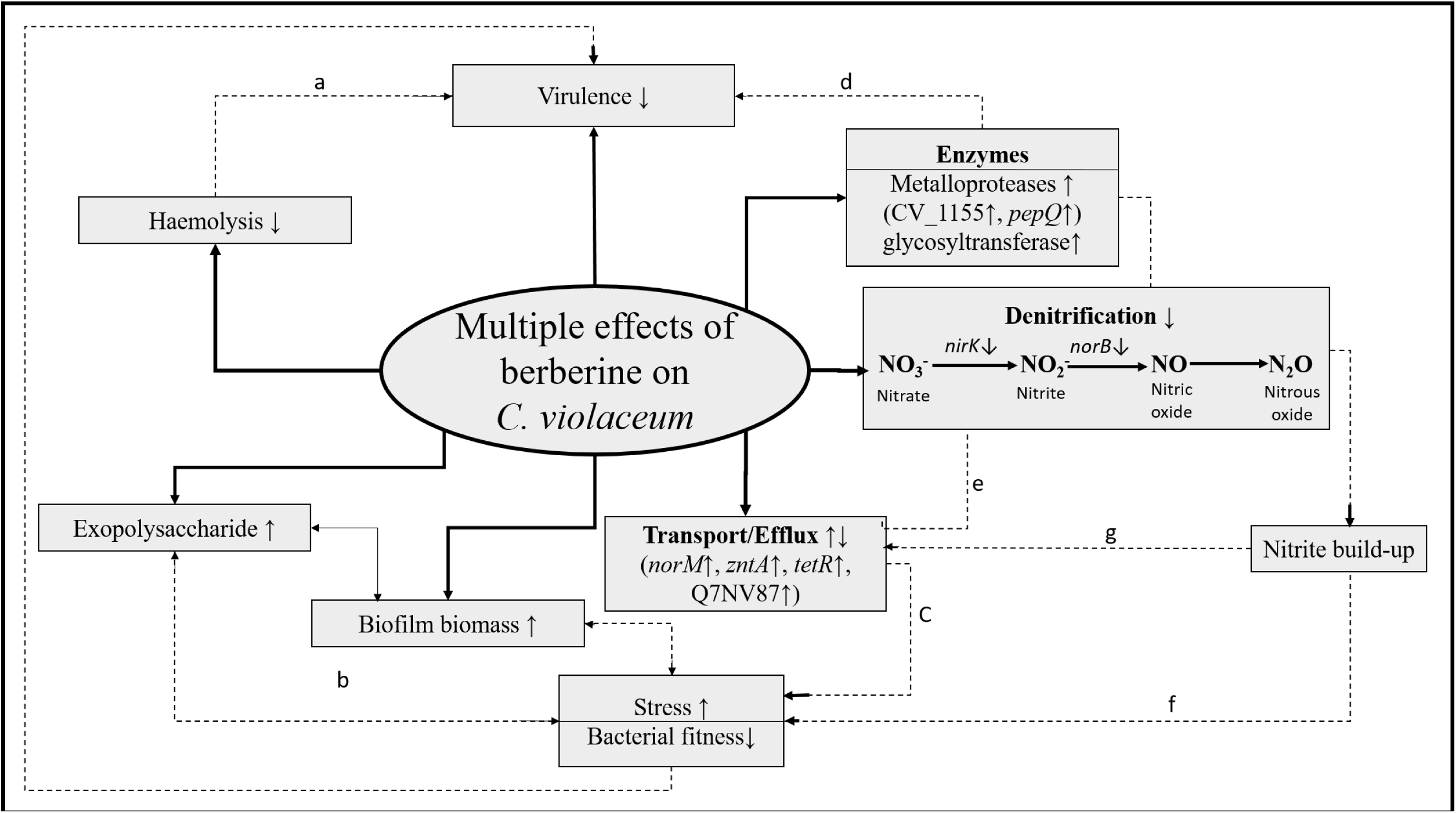
A schematic summary of multiple effects of berberine on *C. violaceum*. (↑) or (↓) arrow indicates the up- or down-regulation of the concerned genes/traits. Solid lines are the directly observed parameters, while dotted lines are interpretations based on literature. (a: Choby and Skaar, 2016; b: Mohammed et al., 2020; c: Sun et al., 2014; d: Miyoshi and Shinoda, 1977; e: Schreiber et al., 2006; f: Bazylinski et al., 1986; g: Grangeiro et al., 2004))

## Supporting information

Legend for supplementary videos

## Acknowledgements

Authors thank NERF (Nirma Education and Research Foundation), Ahmedabad, for infrastructural and financial support, and for providing a doctoral stipend to NT. GG and NT acknowledge support from the Government of Gujarat via their SHODH scheme. VK acknowledges useful discussions with Late Dr. Ashok Vaidya, which contributed towards initiation of this study. N2 Bristol strain was provided by the CGC, which is funded by NIH Office of Research Infrastructure Programs (P40 OD010440).

