## Supplementary figures and tables for "Berberine, the major bioactive compound from *Berberis aristata,* attenuates virulence of multidrug resistant *Chromobacterium violaceum* at non-lethal concentrations by targeting bacterial efflux and denitrification machinery"

### Table S1. Antibiogram of *Chromobacterium violaceum* generated through Kirby-Bauer disc diffusion assay

| Sr. No. | Antibiotic class | Antibiotic | Concentration (µg per disc) | Zone of inhibition (ZOI) | *Chromobacterium violaceum* |
|---|---|---|---|---|---|
| 1 | β-Lactam | Ceftriaxone | 30 | 25±1.41 | S |
| 2 | | Cefpodoxime | 10 | 0±0 | R |
| 3 | | Cefixime | 5 | 0±0 | R |
| 4 | | Cefepime | 30 | 29.5±0.70 | S |
| 5 | | Cefotaxime | 30 | 20.75±1.06 | S |
| 6 | | Ampicillin | 10 | 16.5±2.12 | I |
| 7 | | Ticarcillin | 75 | 24±1.41 | S |
| 8 | | Augmentin | 30 | 0±0 | R |
| 9 | | Imipenem | 10 | 19.5±3.53 | S |
| 10 | Aminoglycoside | Streptomycin | 25 | 25±5.65 | S |
| 11 | | Gentamicin | 10 | 32.5±2.12 | S |
| 12 | | Tobramycin | 10 | 25±2.82 | S |
| 13 | | Amikacin | 30 | 29.5±0.70 | S |
| 14 | | Kanamycin | 30 | 24.5±6.36 | S |
| 15 | Fluoroquinolones/ Quinolones | Ciprofloxacin | 5 | 46.25±0.35 | S |
| 16 | | Levofloxacin | 5 | 47±1.41 | S |
| 17 | | Ofloxacin | 5 | 42±2.82 | S |
| 18 | | Norfloxacin | 10 | 54.25±3.18 | S |
| 19 | | Sparfloxacin | 5 | 41.5±2.12 | S |
| 20 | | Moxifloxacin | 5 | 46.75±1.76 | S |
| 21 | | Gatifloxacin | 5 | 47.75±1.06 | S |
| 22 | | Nalidixic Acid | 30 | 37±4.24 | S |
| 23 | Tetracycline | Tetracycline | 30 | 24±0 | S |
| 24 | | Doxycycline Hydrochloride | 30 | 25.75±2.47 | S |
| 25 | Glycopeptide | Vancomycin | 30 | 13.5±2.12 | I |
| 26 | Polymyxin | Colistin | 10 | 0±0 | R |
| 27 | Sulfonamide | Co-Trimoxazole | 25 | 35.5±6.36 | S |
| 28 | Rifamycin | Rifampicin | 5 | 16±0 | I |
| 29 | Lincosamide | Clindamycin | 2 | 9±1.41 | R |
| 30 | Phenicol | Chloramphenicol | 30 | 27±1.41 | S |
| 31 | Nitrofuran | Nitrofurantoin | 300 | 30±0 | S |

Antibiotic susceptibility profile of the organism was generated using the antibiotic discs- Icosa G-I Minus (HiMedia, Mumbai) through disc diffusion assay on cation-adjusted Mueller-Hinton agar (HiMedia) as per Clinical Laboratory Standards Institute (CLSI) guidelines (https://doi.org/10.1177/001857870403900608). The zones of inhibition were measured and the interpretation (S - sensitive, I - intermediate, R - resistant) was drawn as per zone size interpretative chart provided by the manufacturer.

**Table S2. Quantification of extracted RNA, library, and insert size**

| Sr. No. | Sample name | Quantification of extracted RNA | | Library quantification and insert size analysis | |
|---|---|---|---|---|---|
| | | $OD_{260}/OD_{280}$ | RIN value | ng/µL | Insert size |
| 1 | Control | 2.04 | 8.8 | 24.4 | 296 |
| 2 | Experimental | 2.09 | 8.8 | 22 | 293 |

**Table S3. Time-temperature profile for RT-PCR assay**

| Temperature (°C) | Time (s) | Remarks |
|---|---|---|
| PCR cycles (45 cycles) | | |
| 95 | 15 | Denaturation temperature |
| 59 | 60 | Annealing temperature |
| Melt curve stage | | |
| 95 | 15 | |
| 60 | 60 | |
| 95 | 15 | |

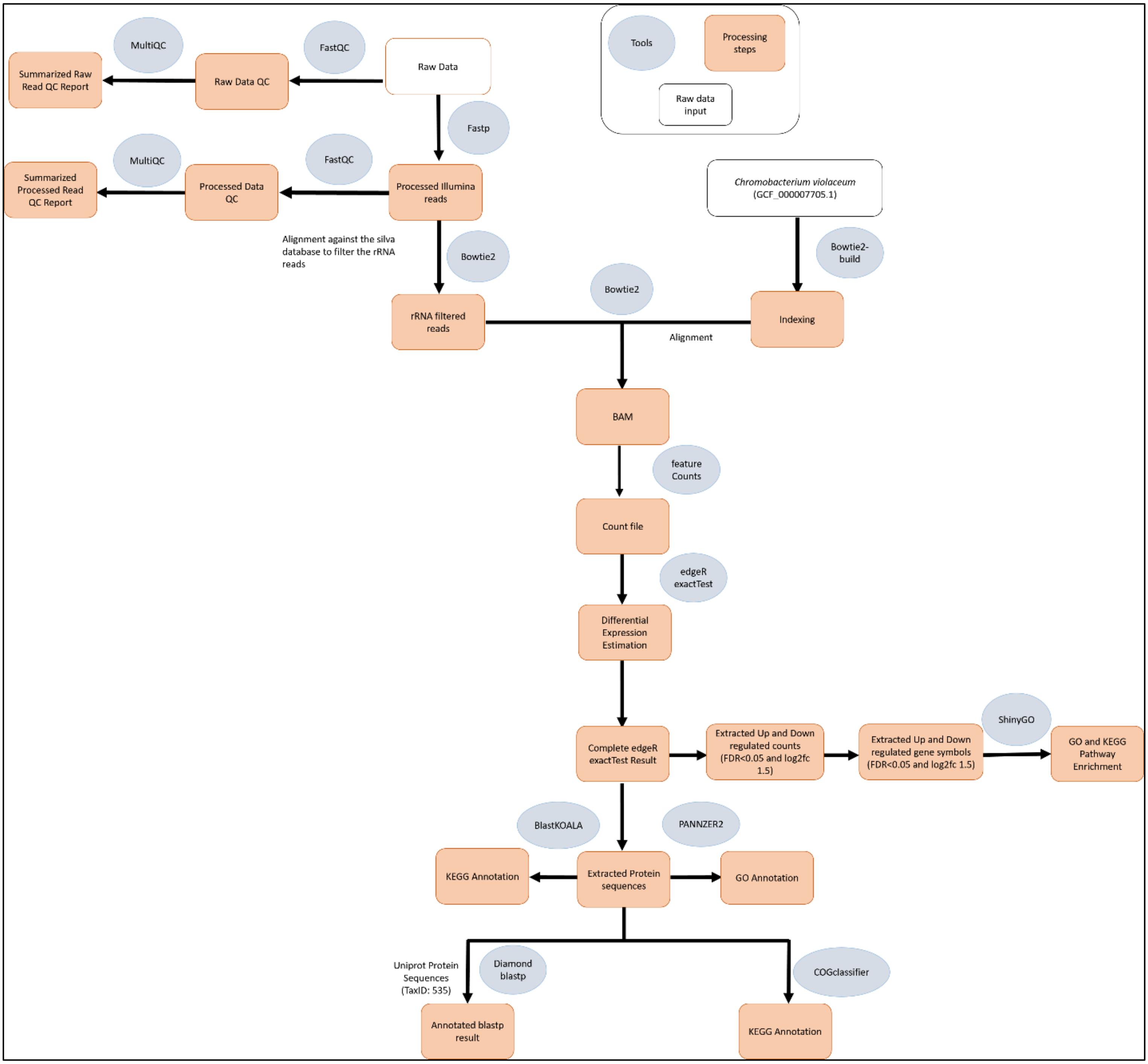


**Figure S1. A schematic presentation of the methodology/workflow employed for whole transcriptome analysis**

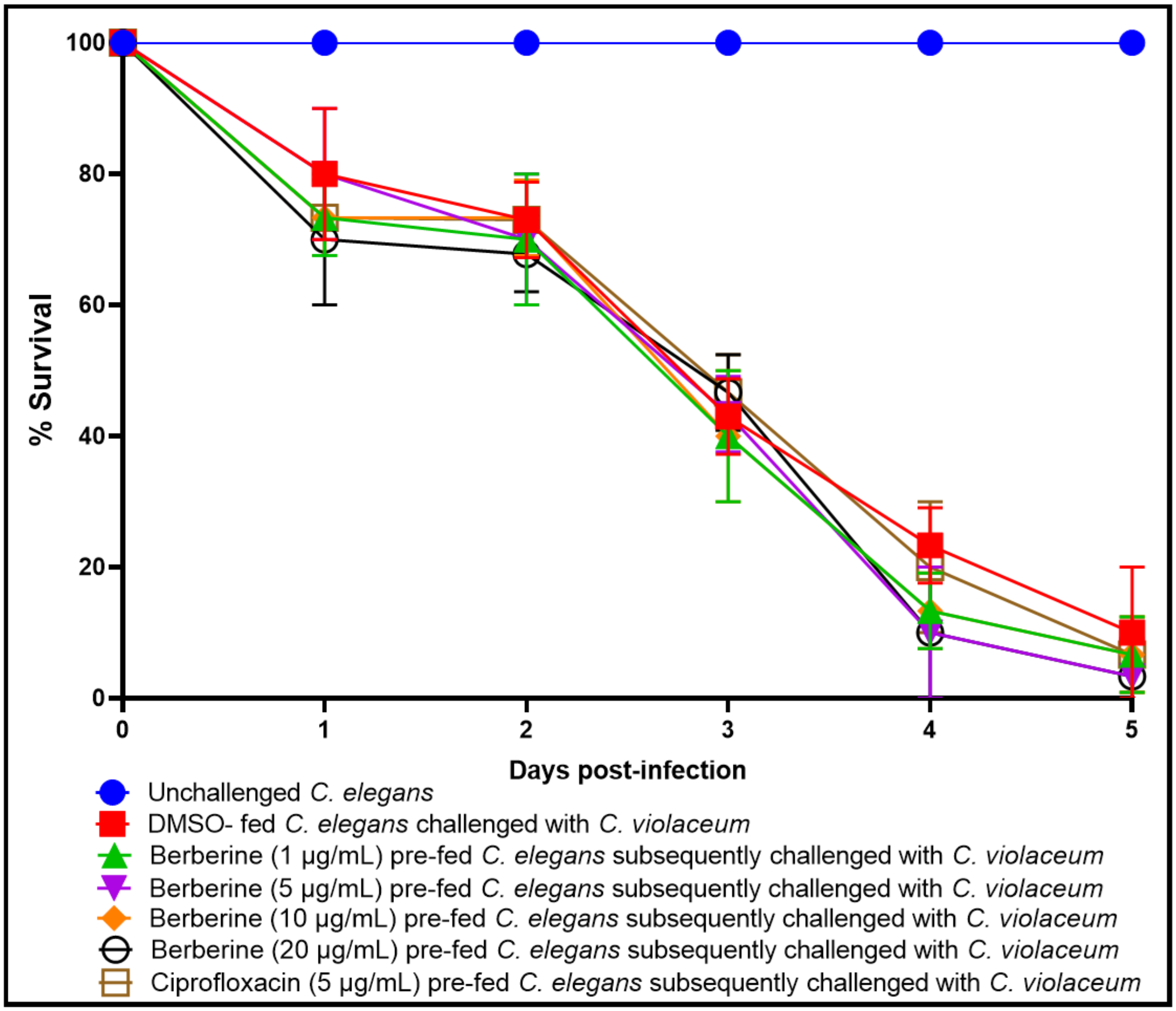


**Figure S2. Berberine did not offer prophylactic protection to the worm population against subsequent pathogen challenge.** Prior feeding (for 72 h) of worms with DMSO (0.5%v/v) or berberine did not alter their susceptibility to subsequent challenge with pathogenic bacteria. DMSO (0.5%v/v) and berberine at tested concentrations showed no toxicity towards the worms. Ciprofloxacin tried as a positive control could also not rescue the worms from subsequent pathogen challenge.

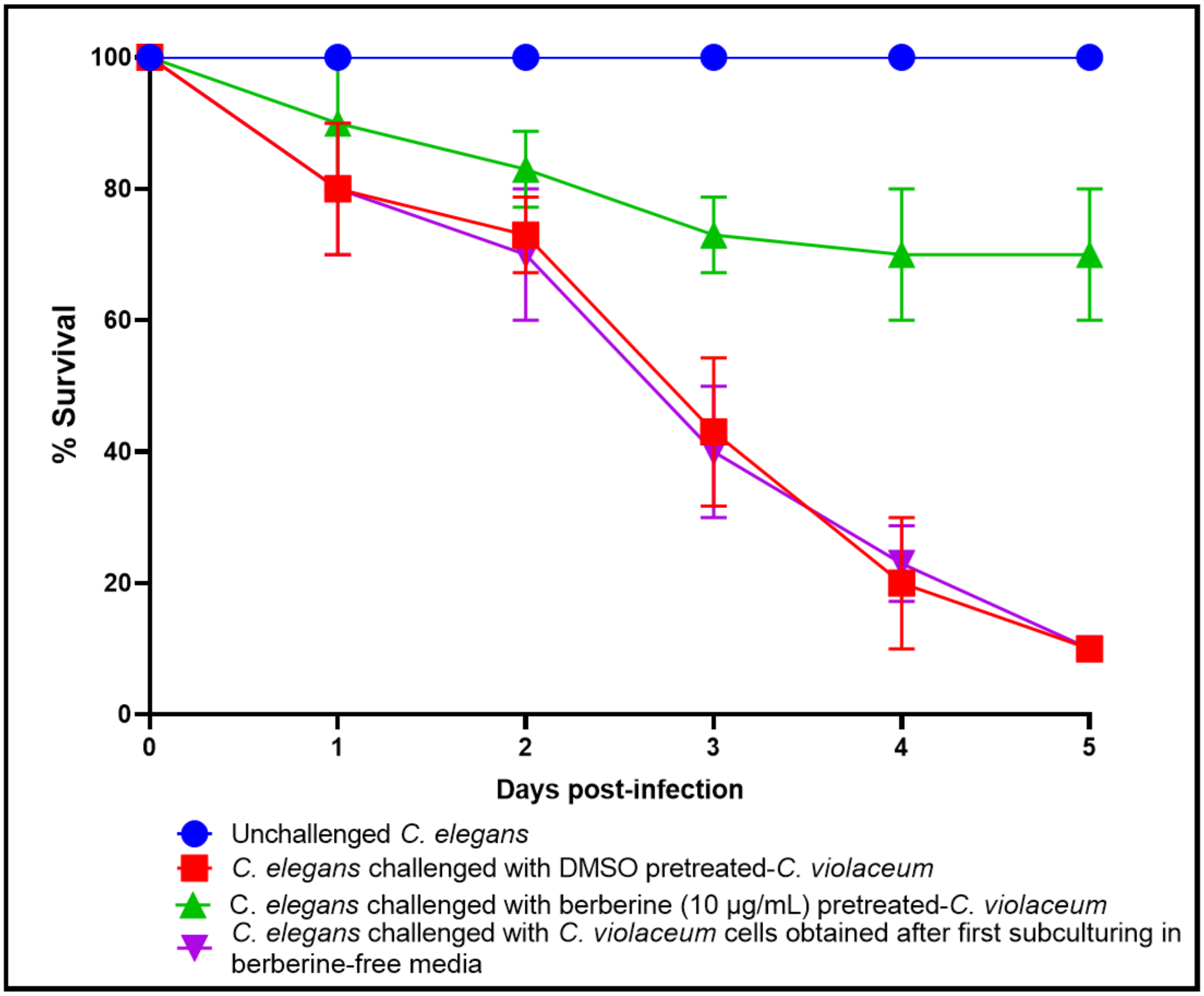


**Figure S3. Berberine had no post-exposure effect on *C. violaceum*.** *C. violaceum* grown in presence of berberine was subcultured in a berberine-free media, and the culture obtained from the latter was compared in terms of its virulence with that of control (no previous berberine exposure) as well as berberine-exposed culture. Berberine's anti-pathogenic effect is not retained upon subculturing of pathogen in berberine-free media.

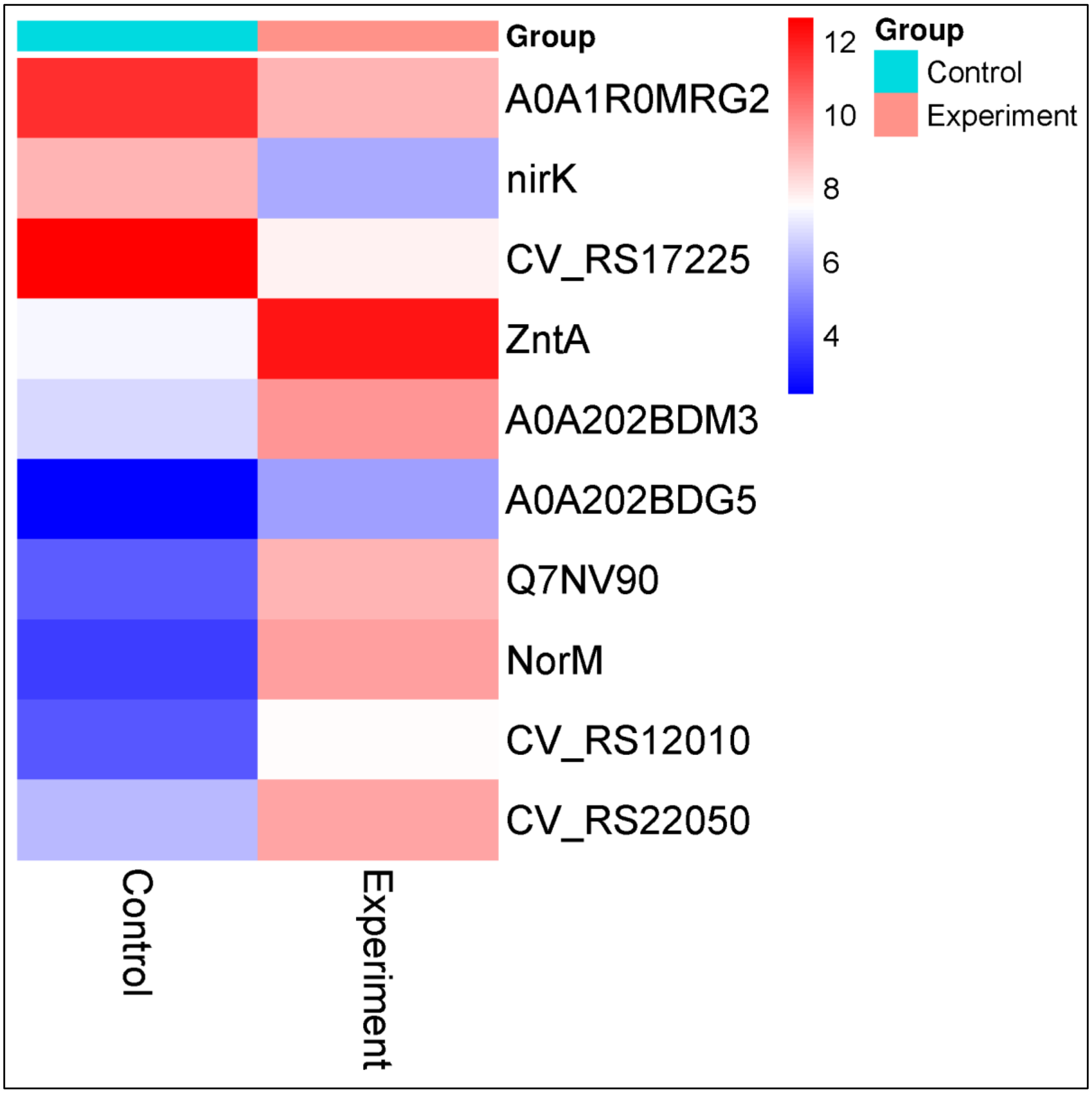


**Figure S4. Heat map of DEGs in berberine-exposed *C. violaceum***

Heat map generated using the online software tool ClustVis (https://biit.cs.ut.ee/clust vis/) showing up-regulated and down-regulated genes with FDR<0.05 and log fold change '±2'.

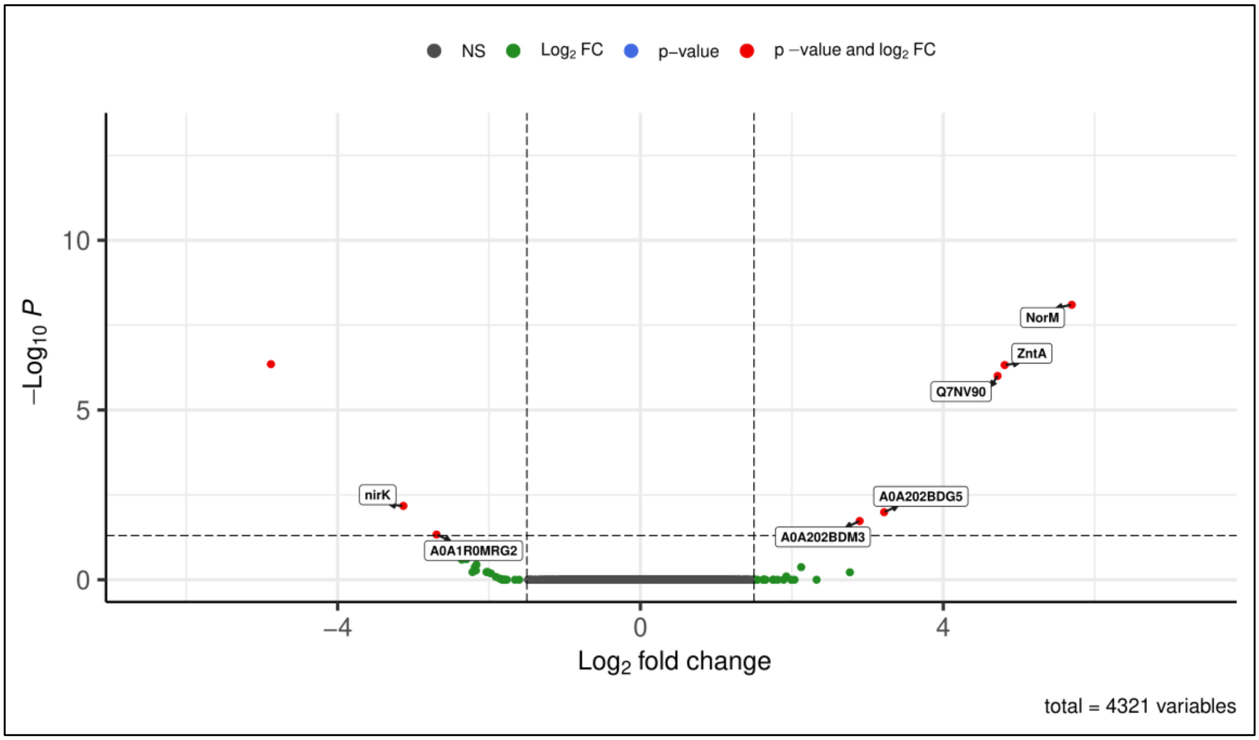


**Figure S5. Volcano Plot of experimental versus control samples**

Volcano plot of expressed genes between the groups Experimental and Control. The y axis illustrates −log 10 p values, and the x axis corresponds to a log 2-fold change of gene expression between Experimental and Control groups. The red points represent statistically significant genes (FDR < 0.05). The genes represented in black are non-significant (NS) and green (log2FC) points denote the ones that were not statistically significant and were not considered for downstream analyses.

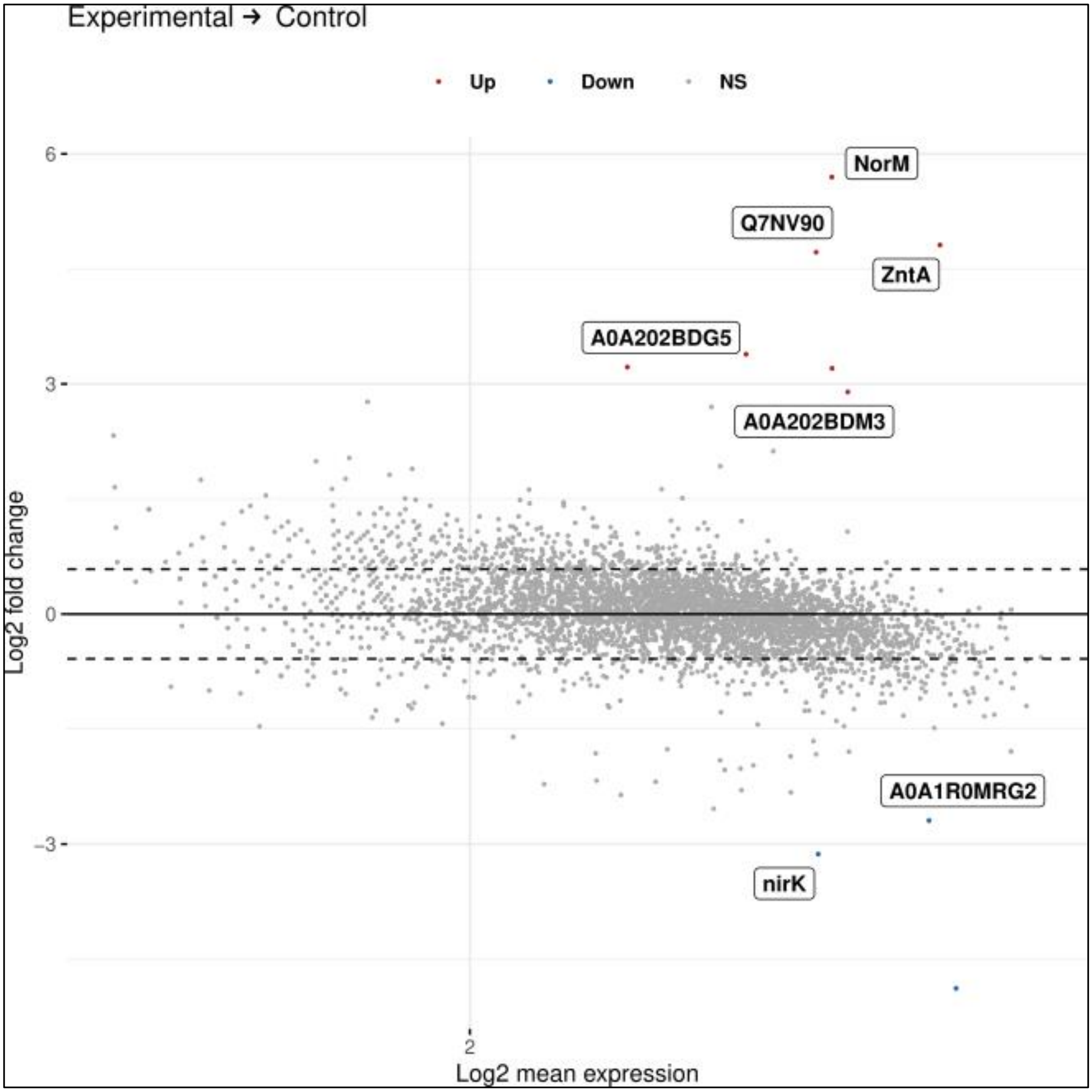


**Figure S6. MA Plot of experimental versus control samples**

The MA-plot shows the distribution of the gene expression between the groups control and experimental. The Y axis shows the Log2fold change (M) and the X axis represents the log of the mean of normalized expression counts (A) of the samples. Red dots correspond to genes up-regulated (> +1.5) and blue dots correspond to genes which are down-regulated (< -1.5) based on the adjusted p-value (FDR<0.05).

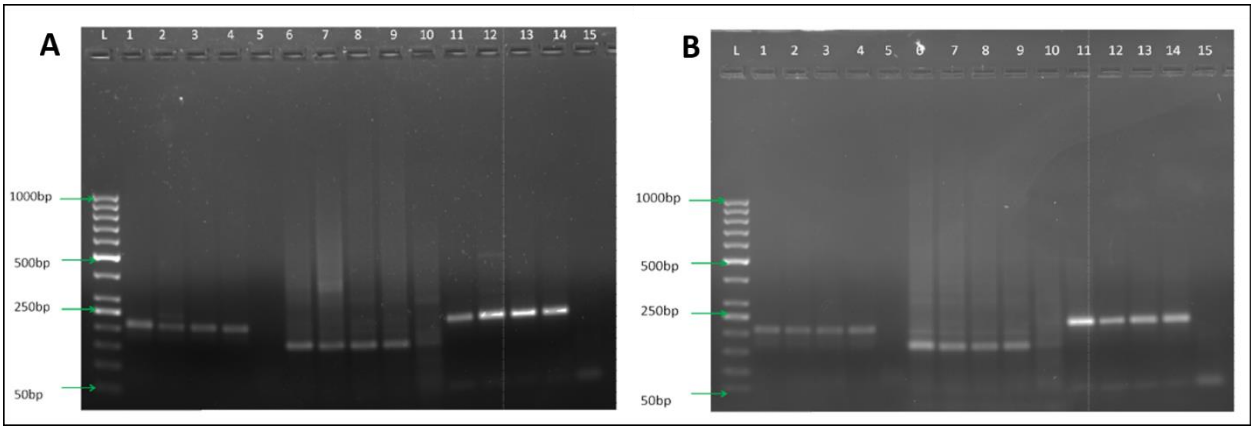


**Figure S7. Gel images corresponding to RT-PCR.** (A) This is from the same nucleic acid sample, which was used for whole transcriptome analysis. (B) This is from an assay independent of the transcriptome assay.1= CV_RS09795 (1/2)_Control; 2= CV_RS09795 (1/2)_Control; 3= CV_RS09795 (1/2)_Experimental; 4= CV_RS09795 (1/2)_Experimental; 5= CV_RS09795 (1/2)_water control; 6= CV_RS17225 (3/4)_Control; 7= CV_RS17225 (3/4)_Control; 8= CV_RS17225 (3/4)_Experimental; 9= CV_RS17225 (3/4)_Experimental; 10= CV_RS17225 (3/4)_water control; 11= CV_RS01880_endogenous_control (5/6)_Control; 12= CV_RS01880_endogenous_control (5/6)_Control;13= CV_RS01880_endogenous_control (5/6)_Experimental;14= CV_RS01880_endogenous_control (5/6)_Experimental; 15= CV_RS01880_endogenous_control (5/6)_Water control
