## Supplementary material for "Berberine, the major bioactive compound from *Berberis aristata,* attenuates virulence of multidrug resistant *Chromobacterium violaceum* at non-lethal concentrations by targeting bacterial efflux and denitrification machinery": Legend for supplementary videos

We investigated anti-pathogenic activity of berberine against a multidrug resistant *Chromobacterium violaceum*, wherein the nematode worm *Caenorhabditis elegans* was employed as a model host for the bacterial pathogen. All these videos were captured at the end of fourth day (i.e. after 96 h incubation), unless specified otherwise. Videos were captured using Magnus Camera (5.1 MP) attached to Labomed Vision 2000 (halogen light source) binocular microscope (4X objective).

**Anti-pathogenic assay:**

*C. violaceum*'s ability to kill the worm is compromised in presence of berberine.

Video A: Control worms in M9 buffer exposed neither to berberine nor bacterial pathogen.

Video B: Vehicle Control: Worms challenged with *C. violaceum* in presence of DMSO (0.5% v/v). DMSO did not interfere with bacterial virulence. Two worms killed by the pathogen are visible.

Video C: Worms challenged with *C. violaceum* in presence of berberine (10 µg/mL) displayed not only higher survival rate and bigger morphology, but they were also able to reproduce (multiple smaller worms are the progenies). Higher agility of surviving worms in this video (as compared to those in video A) indicates them to be in good health.

Video D: Positive Control: Worms challenged with *C. violaceum* in presence of the antibiotic ciprofloxacin (10 µg/mL) displayed higher survival rate. Though antibiotic could rescue them from pathogen-induced death, the surviving worms appear 'sick'.

Video E: Worms in M9 buffer supplemented with berberine (10 µg/ml). Berberine exerted no toxicity towards the worms.

**Anti-virulence (anti-infective) assay:**

*C. violaceum* cells grown in berberine-supplemented media experience an attenuation in their ability to kill the worm host.

Video F: Vehicle Control: DMSO (0.5% v/v)-pre-treated *C. violaceum* could kill 80% of the worms by end of fourth day.

Video G: Worms challenged with berberine (10 μg/mL)-pre-treated *C. violaceum* registered better survival rate. Active movement, bigger morphology, and progeny formation is visible.

Video H: Gentamicin (3 μg/mL) pre-treatment did not attenuate *C. violaceum*'s virulence towards the worm population.

**Post-infection assay:**

Berberine when employed as a post-infection therapy for pre-infected worms, it could save the worm population from pathogen-induced death.

Video I: Vehicle Control: Addition of DMSO (0.5% v/v) post 24-h of pathogen challenge to worms did not interfere with subsequent killing of worms.

Video J: Addition of berberine (10 μg/mL) post 24-h of pathogen challenge to worms not only reduced the death rate, it also supported bigger morphology, more agile movement, and reproduction.

***C. violaceum* exerts cell-mediated killing (and not toxin-mediated) towards the worm population.**

Video K: *C. violaceum* cells suspended in normal saline, when mixed with worm population, kills the worms.

Video L: Supernatant of *C. violaceum* culture fails to exert any toxicity towards the worms. On the contrary, worm population exerts better health and reproduction in presence of the supernatant.

Video M: *C. violaceum* cells suspended in normal saline, when mixed with worm population in presence of berberine, worms exhibit better survival, bigger morphology, and fertility.
